# Insights into *Mus musculus* population structure across Eurasia revealed by whole-genome analysis

**DOI:** 10.1101/2021.02.05.429881

**Authors:** Kazumichi Fujiwara, Yosuke Kawai, Kazuo Moriwaki, Toyoyuki Takada, Toshihiko Shiroishi, Naruya Saitou, Hitoshi Suzuki, Naoki Osada

## Abstract

For more than 100 years, house mice (*Mus musculus*) have been used as a key animal model in biomedical research. House mice are genetically diverse, yet their genetic background at the global level has not been fully understood. Previous studies suggested that they originated in South Asia and diverged into three major subspecies almost simultaneously, approximately 350,000–500,000 years ago; however, they have spread across the world with the migration of modern humans in prehistoric and historic times (∼10,000 years ago to present), and undergone secondary contact, which have complicated the genetic landscape of wild house mice. In this study, we sequenced the whole genomes of 98 wild house mice collected from Eurasia, particularly East Asia, Southeast Asia, and South Asia. We found that although wild house mice consist of three major genetic groups corresponding to the three major subspecies, individuals representing admixture between subspecies are much more ubiquitous than previously recognized. Furthermore, several samples showed an incongruent pattern of genealogies between mitochondrial and autosomal genomes. Using samples likely retaining the original genetic components of subspecies with least admixture, we estimated the pattern and timing of divergence among the subspecies. The results are important for understanding the genetic diversity of wild mice on a global level and the information will be particularly useful in future biomedical and evolutionary studies using laboratory mice established from these wild mice.

## Introduction

The house mouse (*Mus musculus*) has been an important animal model in biomedical research for more than 100 years, and many inbred strains are currently available for research. Inbred laboratory strains are genetically diverse, originating from three wild subspecies (Bonhomme et al. 1987; Didion and Villena 2013; Moriwaki et al. 1984; Yang et al. 2007; Yonekawa et al. 1980, 1981, 1982): North Eurasian *M. musculus musculus* (MUS), South Asian *M. musculus castaneus* (CAS), and West European *M. musculus domesticus* (DOM). The first mouse reference genome sequence was built using a classical inbred strain, C57BL/6J (Chinwalla et al. 2002). Subsequently, dozens of whole-genome sequences of laboratory mouse strains have since been published (Keane et al. 2011). The genome of classical inbred strain is approximately 94.3%, 5.4%, 0.3%, derived from DOM, MUS, and CAS (Keane et al. 2011), respectively, while the mitochondrial genome belongs to DOM (Frazer et al. 2007). Frazer et al. estimated that 10% of the classical inbred strain genome is derived from *M. m. molossinus*, which is thought to have been resulted from hybridization between MUS and CAS (Takada et al. 2013; Yonekawa et al. 1988). In addition, various strains of laboratory mice have been investigated, with some studies analyzing the sequences of diverse mouse strains with a focus on the origin of subspecies (Yang et al. 2011).

Despite intensive effort to sequence the genomes of laboratory mice, genetic diversity of wild house mice has not been thoroughly investigated. Previous studies found that wild house mice are highly genetically diverse and have an estimated effective population size on the order of 10^5^ (Baines and Harr 2007; Geraldes et al. 2008, 2011; Halligan et al. 2010, 2013). Classical inbred strains represent only a small fraction of the genetic diversity of wild house mice (Salcedo et al. 2007). Recent studies revealed a genome-wide pattern of polymorphisms in wild house mice mostly focusing on CAS or DOM within limited geographic ranges (Halligan et al. 2013; Harr et al. 2016; Phifer-Rixey et al. 2018). Therefore, a large-scale genome sequencing study covering the Eurasian continent and surrounding islands would improve our understanding of the worldwide genetic diversity of house mice.

Wild house mice are distributed over most of the world, including remote islands. CAS inhabits the present-day Indian subcontinent and Southeast Asia. DOM inhabits North and South America, Africa, the Middle East, Australia, Southwestern Europe, and many surrounding and remote islands. MUS inhabits Siberia, Central Asia, East Asia, and Northeastern Europe. The homeland of *M. musculus* was proposed to be in the northern part of the Indian subcontinent (Boursot et al. 1993; Din et al. 1996), and its common ancestors diverged into the three subspecies almost simultaneously (Didion and Villena 2013), approximately 350,000–500,000 years ago (Boursot et al. 1996; Geraldes et al. 2008, 2011; Salcedo et al. 2007; Suzuki et al. 2004).

Throughout prehistoric human migration, the wild house mice migrated and lived commensally with humans. With prehistoric and historic long-distance migration, house mice, whose staple foods are grains, expanded their range with the development of agriculture and cultural exchange (Bonhomme et al. 2010; Gabriel et al. 2011; Jones et al. 2013; Moriwaki et al. 1986; Sage 1981). Movement with humans brought genetically diverse subspecies into the secondary contact (Boursot et al. 1993; Duvaux et al. 2011), allowing admixture of their genomes (Bonhomme et al. 2007; Liu et al. 2015; Staubach et al. 2012) despite partial reproductive isolation between subspecies, such as DOM and MUS (White et al. 2011). DOM and MUS come into contact along a narrow hybrid zone in Europe, while CAS and MUS seem to have a broader hybrid zone across Central and Eastern Asia (Boursot et al. 1993; Jing et al. 2014). Previous phylogenetic and phylogeographic studies analyzed mitochondrial DNA sequences and limited nuclear gene sequence data from house mice (e.g., Liu et al. 2008). However, the prevalence of hybridization between subspecies in the global population has not been well studied using genome-wide sequence data. In the pre-genomic era, it was recognized that genetic and phenotypic boundaries between subspecies were obscure due to high variability within subspecies, except the boundary between DOM and MUS in western and central Europe (Boursot et al. 1993). The reason for this obscurity may be that only a limited number of autosomal loci had been analyzed at that time. Moreover, estimating past population size changes allows us to infer the migration history of wild house mice that are commensal to human agricultural culture, which helps the inference of human migration history as well as the origin of subspecies in wild mice.

Rapid advancement in sequencing technologies has enabled the use of whole-genome data to estimate population structures and phylogenetic histories. In this study, we sequenced the whole genomes of 98 wild house mice previously collected from across the Eurasian continent and Southeast Asian islands, focusing on East, Southeast, and South Asian mouse samples covering 16 countries. Our analysis shows a prevalence of hybridization between subspecies beyond the hybrid zones that is greater than previously thought, particularly between CAS and MUS in East Asia. Moreover, we estimated the past population size for all individuals used in this study. Our results provide key understanding of the genetic diversity of house mice at the global level for future biomedical and evolutionary research.

## Results

### Genetic Diversity of Mus musculus

In this study, we analyzed 141 whole-genome sequenced samples of *M. musculus* and *M. spretus*. These included 98 newly sequenced *M. musculus* samples from Eurasia, including those from Estonia (EST), Ukraine (UKR), Russia (RUS), Iran (IRN), Kazakhstan (KAZ), Pakistan (PAK), India (IND), Sri Lanka (LKA), Nepal (NPL), China (CHN), Vietnam (VNM), Indonesia (IDN), Taiwan (TWN), Korea (KOR), and Japan (JPN) (Figure S1). After pruning individuals with equal to or greater than third-degree kinship, which was inferred using the kinship coefficient, 128 samples were used for further analysis (94 of our newly sequenced samples were retained). The list of all samples used in this study is shown in Table S1. For the 94 samples that were newly sequenced, 61 were males and 33 were females. The sex of each of these samples is summarized in Table S2. After the variants were filtered, we obtained 134,030,288 SNVs and 31,618,947 indels for the dataset including 128 samples of *M. spretus* and *musculus*, and 107,337,961 SNVs and 25,875,963 indels for the dataset including 121 samples of *M. musculus*. The overall transition/transversion ratio of our samples was 2.23. The Method section describes the detailed filtering process.

As in Li et al. 2021, mitochondrial genome sequences were clustered into four distinct clades, three of which presumably correspond to CAS, DOM, and MUS subspecies. The mitochondrial genomes of samples from Nepal (NPL01 and NPL02) diverged before the split between CAS and DOM clades. When we classified our newly sequenced 94 samples according to mitochondrial haplogroups, the per-sample nucleotide diversity (heterozygosity) of the three subspecies was in the following ranges: 0.00006–0.00757 for CAS (including NEP mitochondrial haplogroup), 0.00003–0.00450 for MUS, and 0.00040–0.00561 for DOM. The ratio of nonsynonymous to synonymous polymorphic sites for each sample was 0.415 on average. The Table S3 summarizes the basic statistics for each individual, including per-sample nucleotide diversity (heterozygosity) for all our newly sequenced 94 samples.

### Genetic structure of wild house mice

We performed principal component analysis (PCA) by all 128 samples using autosomal 100,832,598 SNVs, including those from *M. spretus* (Figure S2). As shown in Figure S2, all *M. musculus* and *M. spretus* (SPR) were clearly differentiated along principal component 1 (PC1). Principal component 2 (PC2) corresponds to variation within *M. musculus* subspecies. In PC2, *M. musculus* subspecies are differentiated into two clusters with some intermediate samples. Mitochondrial haplotypes and sampling locations indicate that these two clusters largely correspond to the DOM and CAS-MUS groups.

We subsequently excluded SPR and replotted PCA using 84,744,729 autosomal SNVs. Figure 1a shows the locations of samples colored according to the three genetic components identified by the PCA plot (Figure 1b). The PCA eigenvalue for each sample is presented in Table S4. Considering the sampling locations, these three clusters correspond to the CAS, DOM, and MUS subspecies, whereas a wide range of admixture between CAS and MUS clusters was observed (Figure 1b). In the PCA plot, Nepalese samples with distinct mitochondrial haplogroups were clustered with CAS samples. PC1 shows the genetic differences between CAS and DOM, while PC2 shows the genetic differences between CAS and MUS. As shown in the Figure 1a and 1b, the prevalence of hybrid individuals is evident, particularly between CAS and MUS. The Chinese samples are widely scattered throughout a wide range of the CAS-MUS cline. The Japanese samples were also scattered along the CAS-MUS cline, although the range was narrower than in Chinese samples and skewed toward the MUS cluster. We also performed PCA using only X-chromosomal SNVs (Figure S3). This pattern was the same as that using autosomal SNVs; however, the plots were more tightly clustered at each vertex, indicating that admixture was less pronounced on the X chromosomes. For the X chromosome, the PCA eigenvalue for each sample is also shown in Table S4.

**Figure 1.**
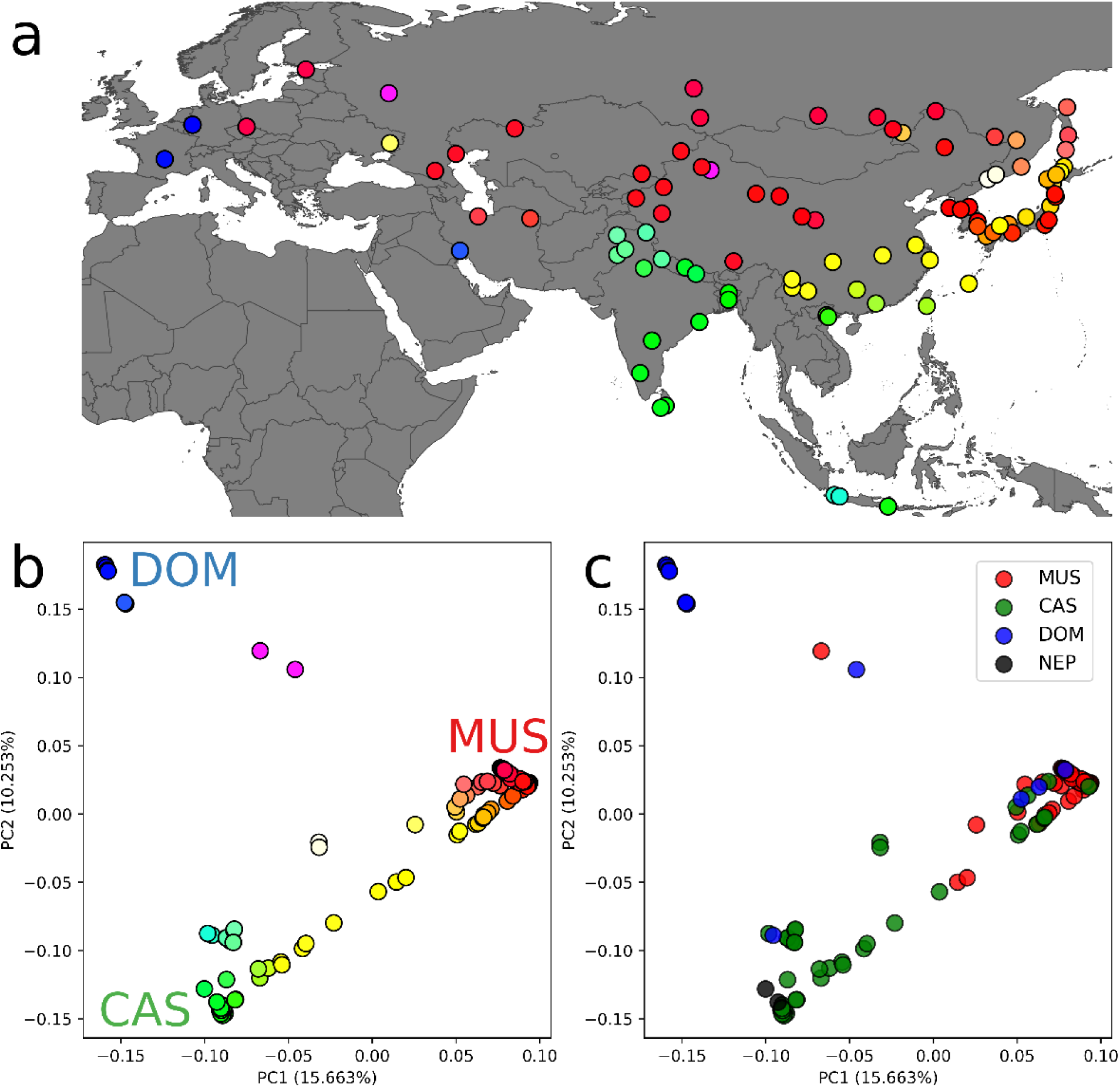
PCA results in wild house mice. (a) A geographical map of sampling locations. The circle represents each individual and the assigned color is the same as the color in panel b. Figure S1 shows the detailed names of sample collection sites. (b) PCA plot of wild house mice using autosomal SNVs. The *x* and *y* axes represent the eigenvalues of PC1 and PC2, respectively. The circles are colored according to “Maxwell’s color triangle,” assigning three vertices to the RGB colors. The red, green, and blue color intensities correspond to MUS, CAS, and DOM genetic components, respectively. The proportion of variance for each eigenvalue is shown in parentheses on the axis label. (c) PCA plot of wild house mice using autosomal SNVs labeled with the mitochondrial genome haplogroup of each sample. The proportion of variance for each eigenvalue is shown in parentheses on the axis label.

In Figure 1b, samples from Germany, Bangladesh, and Korea are located at the vertices of the triangle (Table S4). However, this pattern does not necessarily imply that these samples are representatives of each of the three major subspecies. The pattern may rather be generated by the strong genetic drift in the three populations. In order to identify samples that represented the smallest amount of gene flow between subspecies, we computed *f*_3_ and *f*_4_ statistics among different individuals (Tables S5–S7 and Tables S8–S13, respectively). We found that Indian CAS samples from mountainous regions, western European DOM samples, and Korean MUS samples showed the smallest amount of gene flow with other subspecies. Therefore, we selected an Indian sample (IND04), a Korean sample (KOR01), and a German sample (DEU01) as our reference samples of CAS, MUS, and DOM subspecies, respectively. We also found that one sample collected from Kathmandu in Nepal (NPL02) was the most distantly related to CAS, MUS, and DOM samples. The other sample from Nepal (NPL01) showed a similar pattern, but NPL01 was more closely related to CAS than NPL02 was.

Except for these samples, most CAS-MUS samples exhibited some extent of admixture between subspecies. For example, an Indian sample from Delhi (IND02) was slightly genetically closer to MUS (KOR01) than IND04. The Z score of *f*_4_ (SPR, KOR01; IND04, IND02) was 2.814 (Table S14). Likewise, other CAS samples from neighboring regions, such as Pakistan and Bangladesh, were more similar to MUS (KOR01) than IND03, IND04, and IND07 individuals (Tables S11 and S12). CAS samples from East and Southeast Asian regions also showed various levels of admixture with MUS genomes. Likewise, all MUS samples in East Europe showed a significantly closer relationship to DOM than the other Asian MUS samples. For example, the Z scores of *f*_4_ (SPR, DEU01; KOR01, CZE01) and *f*_4_ (SPR, DEU01; KOR01, KAZ01) were 14.724 and 11.104, respectively (Table S11). Majority of MUS samples in East Asia, particularly samples from Northern China and Japan, showed a high level of admixture with CAS. Among DOM samples, those from outside of West Europe, such as Iranian and Russian samples, showed slightly but significantly closer affinity to MUS and CAS samples than the western European samples did (Tables S10 and S11).

The PCA plot inferred using nuclear genome data was labeled with the four mitochondrial haplogroups (CAS, DOM, MUS, and NEP; Table S3) and is shown in Figure 1c. Most individuals showed congruent patterns in mitochondrial and nuclear genomes, but some samples showed incongruent patterns. Particularly, in the PCA plot (Figure 1b and 1c), the mitochondria–nuclear incongruence was more commonly observed in the MUS cluster than in the CAS or DOM cluster. For example, a sample from Khabarovsk in the Russian Far East (RUS13) had a DOM-type mitochondrial genome but a nuclear genome that was highly similar to MUS.

We also performed an ADMIXTURE analysis to infer the genomic ancestries of *M. musculus* genomes (Figure 2). We analyzed 121 *M. musculus* samples by selecting *K* values from 1 to 5, where *K* is a predefined number of ancient components. We also performed cross-validation to infer the most suitable *K* value (Figure S4), and we found that the cross-validation error rate was the lowest at *K* = 4. At *K* = 3, we confirmed the presence of three genetic components corresponding to CAS, DOM, and MUS. Most Japanese and Chinese samples showed a hybrid pattern with CAS and MUS components. At *K* = 4, another genetic component represented the specific genetic features of Japanese and Korean mice. This component corresponds to *M. musculus molossinus*. The ADMIXTURE results using X-chromosomal SNVs were essentially the same as those using autosomal SNVs (Figure S5).

**Figure 2.**
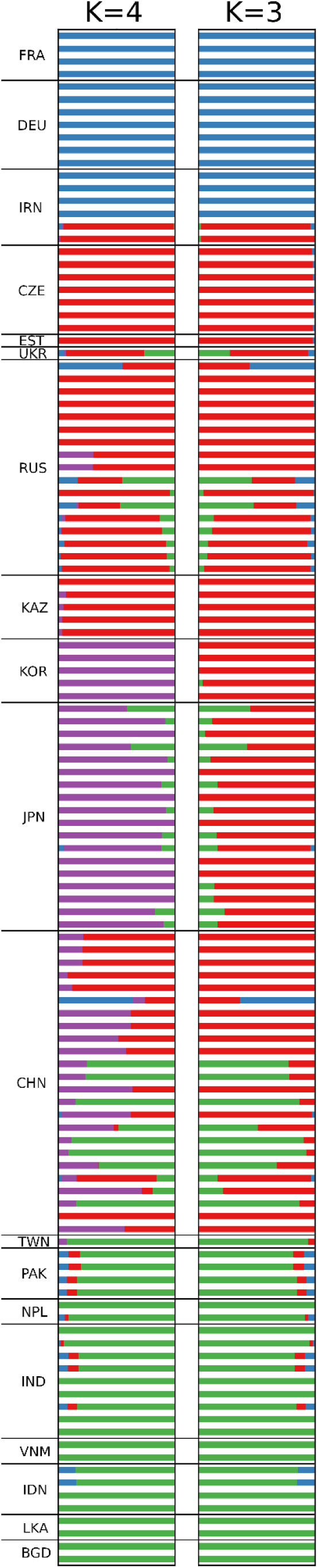
ADMIXTURE plot of autosomes showing the proportion of estimated subspecies genetic components. Results for cluster *K* = 3 and *K* = 4 are presented. Samples with the same country code are ordered according to sampling site from west to east and north to south.

### Inference of past demography using PSMC and MSMC

To estimate the past demographic pattern of wild house mice, we performed Pairwise Sequentially Markovian Coalescent (PSMC) analysis for each individual. Figure S6 presents the PSMC plots of all individual samples. Although some of the samples showed unusual patterns, the trajectories of most samples were largely classified into three categories, each representing CAS, DOM, and MUS (Figure S6). Each subspecies experienced a similar population history up to about 100,000 years ago, but after that, we observed components that experienced different population histories at each of the sampled sites. In particular, the MUS and DOM showed fluctuations in population size due to the bottleneck effect.

In order to further investigate the past demography in more recent times, we also performed Multiple Sequentially Markovian Coalescent (PSMC) analysis using MSMC/MSMC2 software, which utilized the information of multiple haplotypes from each population. In Figure 3, we show the results of four haplotypes from two Indian samples (IND03, IND04) for CAS, eight haplotypes from four German samples for DOM (DEU01, DEU03, DEU04, and DEU06), and eight haplotypes from four Korean samples for MUS (KOR01– 03 and 05). Note that, because the number of analyzed haplotypes was smaller in CAS than in DOM and MUS, the demography of CAS in recent days (after ∼10,000 years ago) may not be estimated reliably. We assumed a mutation rate of 5.7 × 10^−9^ per base pair per generation (Milholland et al. 2017) for converting generation number to years. Three subspecies followed different trajectories of population size change. The effective population size of CAS increased in ancient time period around 100,000 years ago and later continued to decrease. After the decline, the effective population size of CAS reached a maximum around 1,000–3,000 years ago. In contrast, both MUS and DOM experienced population shrinkage after 300,000–500,000 years ago. The trajectories of population sizes in DOM and MUS were very similar prior to 100,000 years ago. DOM later experienced two rounds of population size expansion, around 50,000 and 5,000 years ago. Although MUS experienced small population expansion events, their population size continued to decrease after 500,000 years ago. The trajectories of Korean MUS showed the signature of recent population bottleneck and expansion, probably between 2,000–4,000 years ago. DOM also seems to have experienced recent population bottleneck and expansion between 4,000–6,000 years ago.

**Figure 3.**
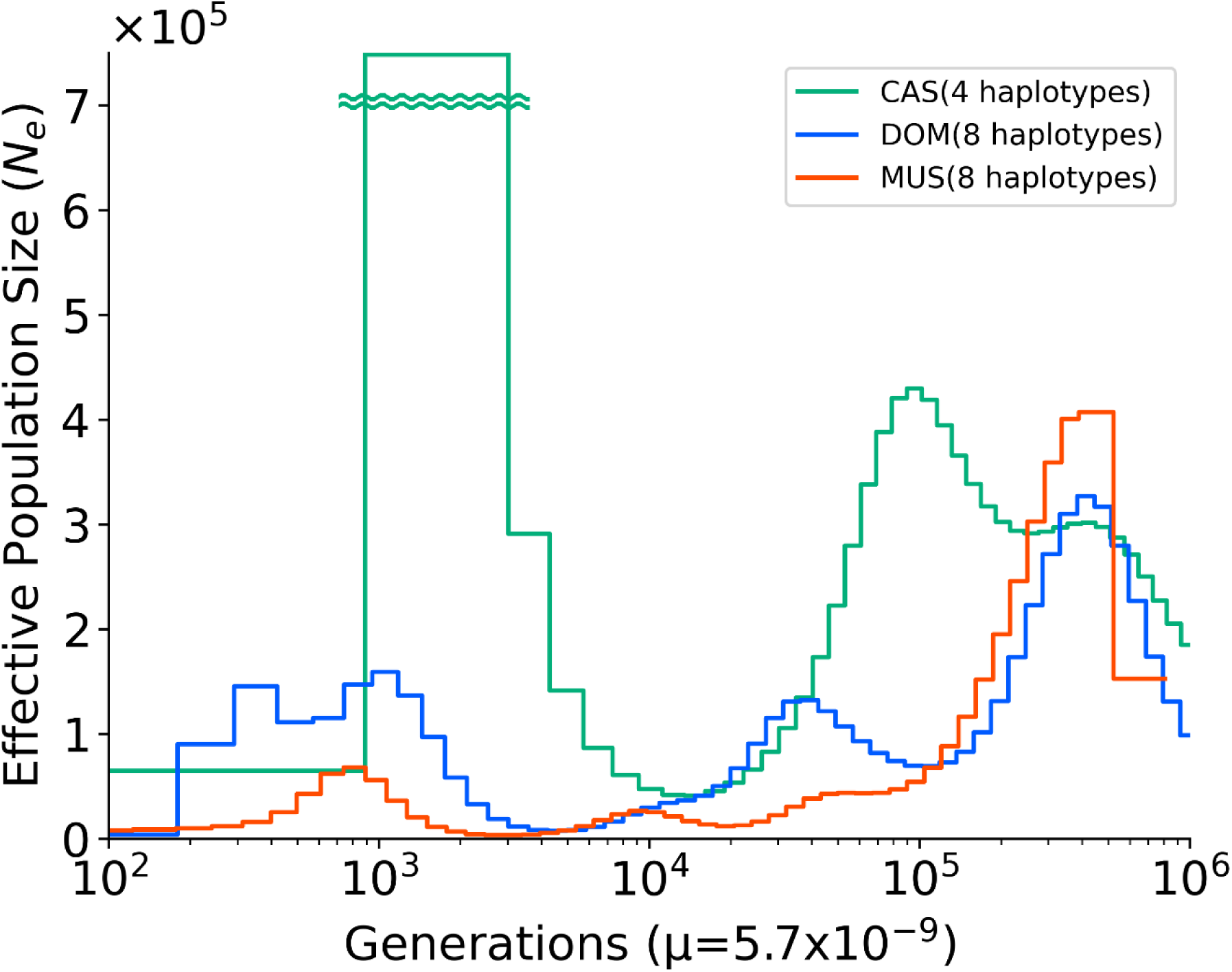
Inferred population sizes as determined using MSMC analysis. The *x* axis represents time before present scaled by the mutation rate of 0.57 × 10^−8^ per site per generation, and the *y* axis represents the effective population size. The red, green, and blue lines represent past population sizes of Korean (MUS), Indian (CAS), German (DOM) samples, respectively. Areas that are far out of the display range are represented by wave lines.

### Genetic relationship among three subspecies

Previous phylogenetic studies using partial or complete mitochondrial genome sequences showed a sister relationship between CAS-DOM (Li et al. 2021; Geraldes et al. 2008) and CAS-MUS (Jing et al. 2014), while genome sequencing studies of wild-derived mice suggested a CAS-MUS clade (White et al. 2009; Keane et al. 2011; Phifer-Rixey et al. 2020). First, we performed *f*-statistics analysis using representative samples of each subspecies that is, IND04 for CAS, DEU01 for DOM, and KOR01 for MUS. We calculated outgroup *f*_3_ statistics for all pairs among the three target samples, making SPR an outgroup (Figure 4). A larger *f*_3_ statistic value indicates that the two subspecies share a larger amount of genetic drift, which means a closer relationship between the two subspecies. The CAS-MUS pair showed an *f*_3_ value that was statistically larger than the other comparisons between subspecies (*p* = 3.12 × 10^−8^ between CAS-MUS and CAS-DOM, *p =* 6.22 × 10^−7^ between CAS-MUS and DOM-MUS; Tukey’s test). To confirm this result, we performed four population tests using *f*_4_ statistics. The Z scores for *f*_4_ (SPR, DEU01; IND04, KOR01), *f*_4_ (SPR, IND04; DEU01, KOR01), and *f*_*4*_ (SPR, KOR01; DEU01, IND04) values were 2.003, 28.518, and 16.001, respectively (Tables S10, S8, and S13). These results support a close genetic relationship between CAS and MUS. Tables S15–S17 and Tables S18–S23 present the *f*_3_ statistics and four *f*_4_ statistics for chromosome X, respectively.

**Figure 4.**
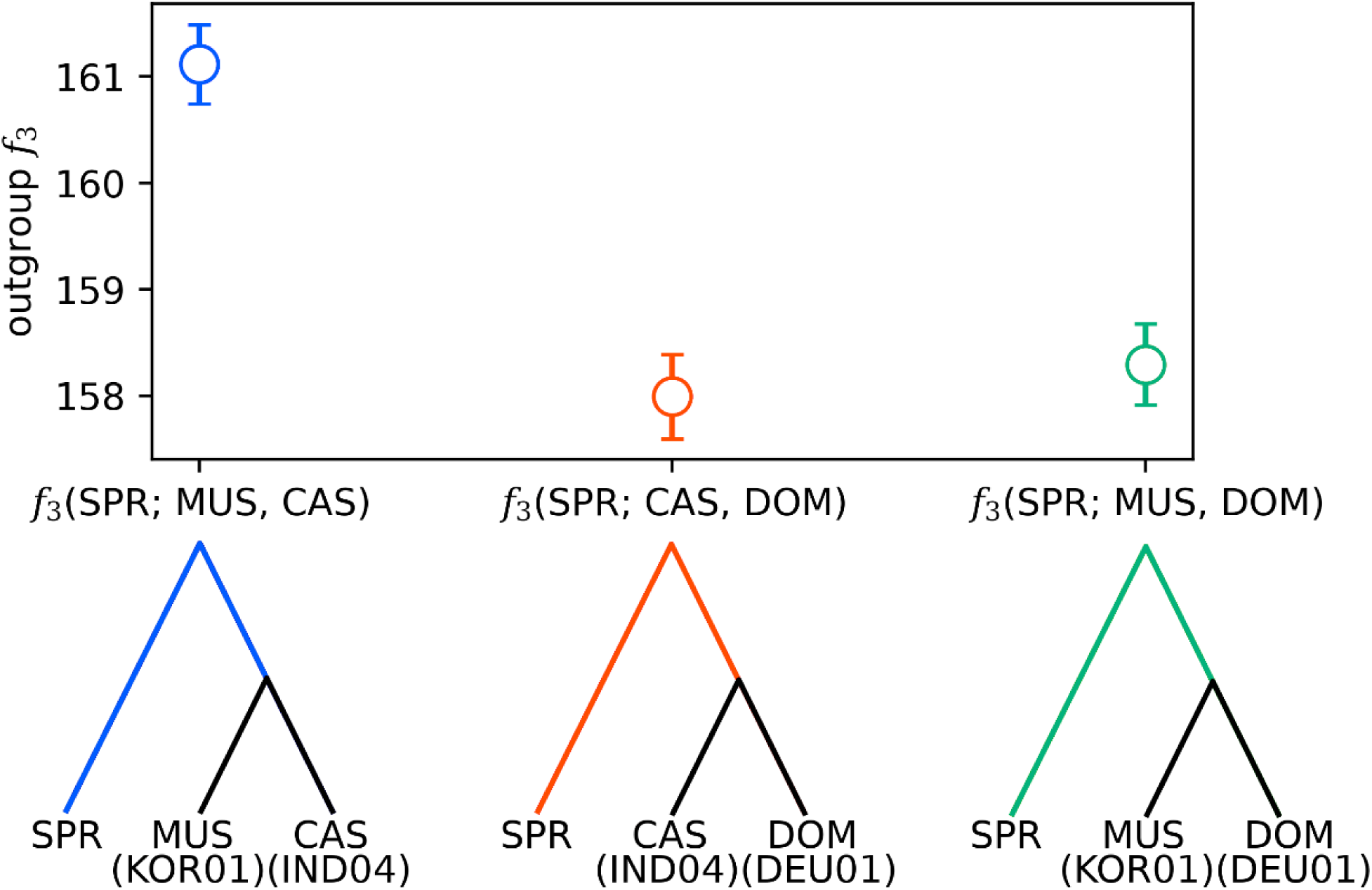
Genetic distances between subspecies using outgroup *f*_3_ statistics. The *y* axis indicates the outgroup *f*_3_ statistics using SPR as the outgroup. Higher *f*_3_ values indicate that the two target populations shared more genetic drift, implying that the two populations diverged more recently or experienced a larger amount of gene flow between populations.

We subsequently constructed a neighbor-joining tree using pairwise genetic distances between samples, initially including all samples, and SPR as the outgroup (Figures S7 and S8). The tree presented a sister relationship between DOM and MUS with exceptionally longer blanch of DOM clade, which is contradictory to the *f*_3_ and *f*_4_ statistics. However, we found that two particular samples, CHN06 from Urumqi and RUS01 from Moscow, which showed the strong signature of hybridization between MUS and DOM, are potentially distorted the pattern. To solve the problem, we reconstructed a tree excluding all potential hybrid individuals. We reconstructed a phylogenetic tree using only individuals with more than 80% signal of one subspecies ancestry from ADMIXTURE results and obtained the pattern of CAS-MUS clade (Figure 5).

**Figure 5.**
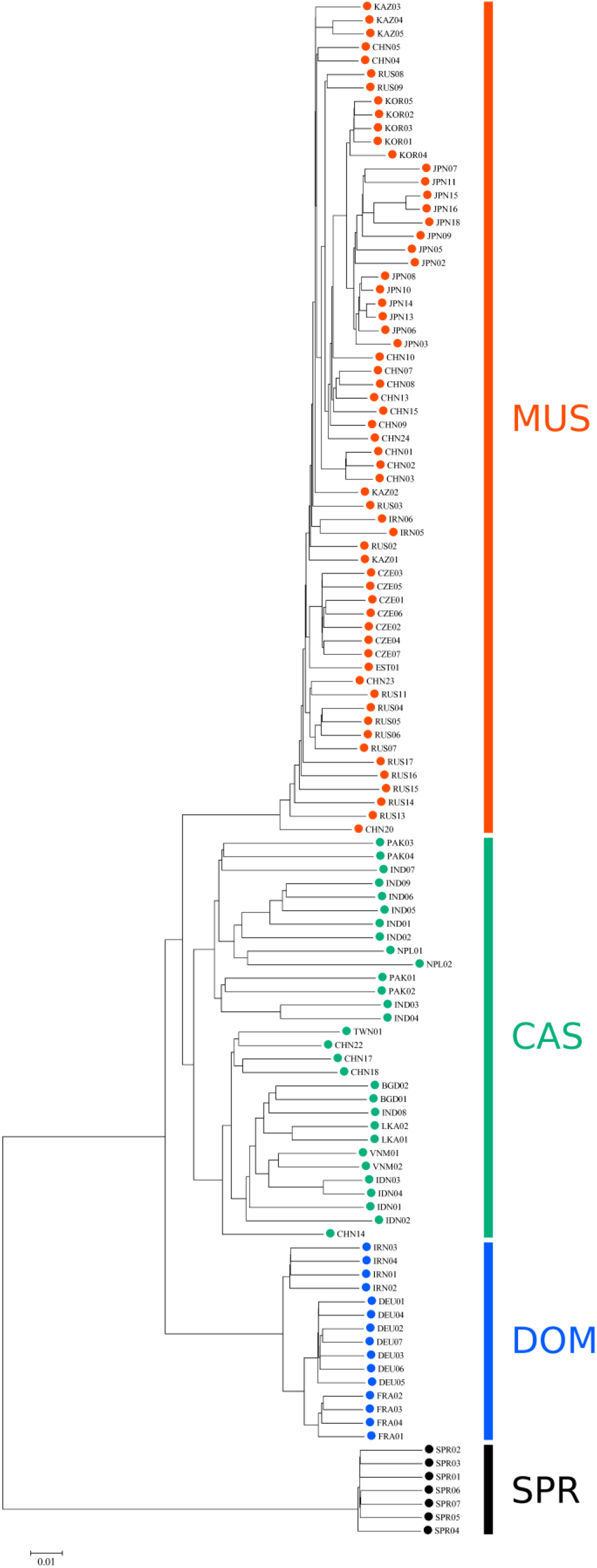
A phylogenetic tree showing the genetic relationships of the three major subspecies, reconstructed by excluding hybrid individuals. The label colors of red, green, and blue are corresponds to the MUS, CAS, and Dom, respectively. The *M. spretus* is used for the outgroup of the *M. musculus*.

### Estimation of divergence times between subspecies

In order to estimate the divergence timing of three subspecies, we performed the cross-subspecific MSMC analysis (Figure 6). We initially used Indian, German, and Korean samples and as the representative samples for each subspecies but found that the strong population bottleneck that occurred in the ancestors of Korean samples made it difficult to correctly infer the population history before the bottleneck. We, therefore, used Kazakhstan samples as representatives of MUS. Although Kazakhstan samples showed some level of admixture with DOM, it would not affect the estimated subspecies divergence time if the admixture occurred more recently (after 10,000 years ago). The divergence times between CAS and DOM, CAS and MUS, DOM and MUS were separately estimated using the time point when the relative cross-coalescent rate (rCCR) equals to 0.5. We found that the divergence between CAS and MUS was the most recent (95% CI:187,365–188,647), consistent with the above analyses. The divergence time between CAS and DOM and DOM and MUS were nearly the same: 223,614–225,306 and 245,411–247,175 years ago (95% CI), respectively.

**Figure 6.**
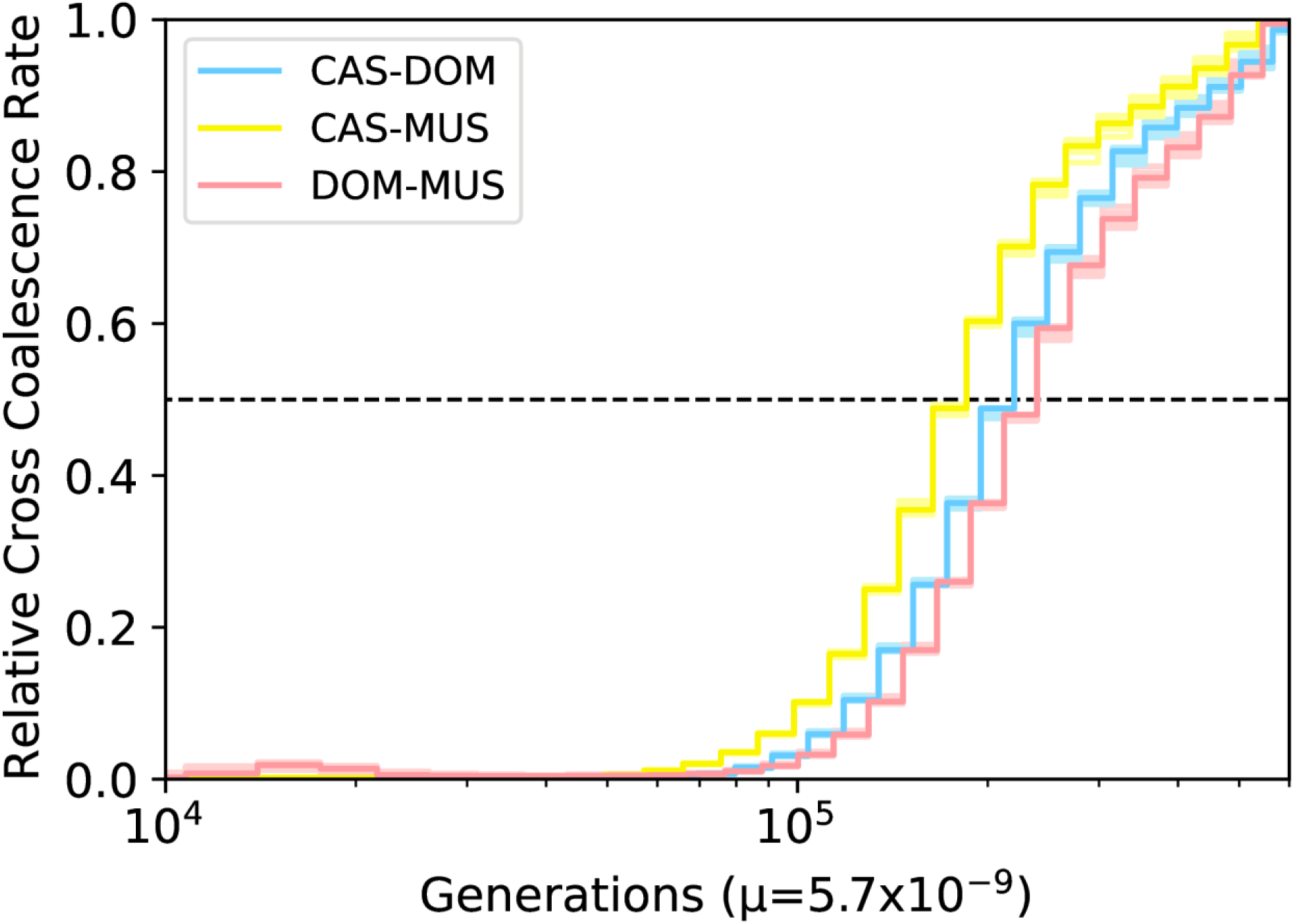
Diagram of the divergence process of *M. musculus* subspecies by cross-coalescence rate demonstrated by the Multiple Sequentially Markovian Coalescent (MSMC) method. The *x* axis represents time before present scaled by the mutation rate of 0.57 × 10^−8^ per site per generation, and the *y* axis represents the relative cross coalescence rates (rCCR). The Magenta, Cyan, Yellow lines correspond to the DOM-MUS, CAS-DOM, and CAS-MUS relative cross coalescence rates change. The dotted line shows the rCCR=0.5 point, which is heuristically identified as the estimated time of split time between the two populations.

## Discussion

Although house mice have been widely used in biomedical research, the global genetic landscape of wild house mice has not been clear. Previous studies analyzed genome sequences focusing on West European (DOM), Middle Eastern (CAS), and North American (DOM) samples (Harr et al. 2016; Mack et al. 2018; Staubach et al. 2012). This is the first genome-wide study of wild house mice including three subspecies that focuses on genetic diversity across the Eurasian continent and Southeast Asian islands. This study clarifies the present and ancestral population structure of the species. Wild house mice have a much larger effective population size than the human population (Geraldes et al. 2008, 2011; Halligan et al. 2010, 2013), and our data confirmed that the nucleotide diversity of wild house mice is much higher than that of humans. Based on the results of the phylogenetic tree excluding the hybrid individuals shown in Figure 5, we calculated the nucleotide diversity in each subspecies; the nucleotide diversity of wild house mouse was 0.527% for CAS, 0.244% for DOM, and 0.225% for MUS. Considering that the average human nucleotide diversity is ranged from 0.08% to 0.12%, we confirmed that the nucleotide diversity of wild house mouse is much higher than that in humans (Arbiza et al. 2014; Prado-Martinez et al. 2013; Perry et al. 2012). These values are consistent with previous studies (Harr et al. 2016; Geraldes et al. 2008; Phifer-Rixey et al. 2014), which showed that CAS has the highest diversity in all subspecies.

PCA results in Figure 1b show a wide spectrum of samples within CAS and MUS genetic clines. In particular, the Japanese and Chinese samples exhibit a wide range of distribution along the PCA plot. Even samples from other locations, such as Southeast Asia, showed an admixture signature to some extent. ADMIXTURE results (Figure 2) largely agreed with PCA results. Our analysis using *f*_3_ and *f*_4_ statistics showed that samples from India, Germany, and Korea had the smallest genetic component derived from different subspecies. However, these results do not necessarily mean that the samples represent genuine subspecies with pure ancestry and without admixture between subspecies. This could be clarified by further sampling of wild mice in the Eurasian continent.

The PSMC results presented here indicate that each of the three subspecies has experienced similar population size changes within the subspecies (Figure 3), but that each individual has been undergone a different history in relatively recent years. As an example of MUS, comparing CHN03 (China: Aksu), EST01 (Estonia: Talinn), and IRN06 (Iran: Mashhad), all of them show similar population size changes until ∼50,000 years ago, but more recently, the Chinese Aksu population experienced a strong bottleneck for 20,000–30,000 years ago and then experienced an increase in population size about 10,000 years ago (Figure S6). The population increase about 10,000 years ago was also observed in the Iranian Mashhad population, but it did not show the extreme bottleneck like in the Chinese Aksu population. In contrast, the Estonian Talinn population did not show any increase in population size around 10,000 years ago, suggesting that these populations experienced distinctly different population size changes depending on the regions. Furthermore, as mentioned above, wild house mice in certain regions, such as in individuals on Russia, Japanese archipelago and Korean peninsula, are subject to the extreme population bottleneck, and it makes difficult for some individuals to follow past genetic demography due to the large loss of polymorphic markers. For example, we were not able to trace their population history before 100,000 years ago due to the strong bottleneck effect (Figure S6) for the RUS06 (Russia: Irkutsk), JPN13 (Japan: Ashiro), and KOR01 (Korea: Baengnyeong Island) samples.

Our results may substantially change the simple trinity view of *M. musculus* subspecies. The observed pattern implies that admixture between CAS and MUS has continued since 10,000 years ago in Asia and that many “MUS-like CAS” and “CAS-like MUS” samples exist. This complex pattern has not been captured by mitochondrial phylogeny. Indeed, we observed many cases with incongruence between mitochondrial genotypes and autosomal genotypes. In particular, many Japanese samples harbor CAS-type mitochondrial genomes and highly MUS-like nuclear genomes (Figure 1c). Li et al. (2021)’s study of whole mitochondrial genome sequences suggested that the CAS-type mitochondrial genome was introduced to the Japanese archipelago in the late Neolithic period (∼3800–3200 years ago) and that the MUS-type mitochondrial genome migrated later (∼2700 years ago) and quickly spread to the archipelago. The latter migration of house mice may coincide with the introduction of intense rice farming to the Japanese archipelago (Li et al. 2021). Such a pattern of nuclear–mitochondrial genotype mismatching was also reported in New Zealand house mice (Veale et al. 2018). Mechanisms that explain the incongruence between mitochondrial and nuclear genotypes are largely unknown; however, sex-biased migration and/or natural selection may be underlying mechanisms.

The analysis of past demography revealed different trajectories of population sizes in three subspecies. MSMC results showed that the population size decline of DOM and MUS started around 300,000–400,000 years ago. The timing corresponds to the beginning of rCCR decline between three subspecies (Figure 6) and may be associated with the process of population divergence. Interestingly, both DOM and MUS experienced recent strong population bottleneck and expansion, which would be associated with the spread of agriculture. The entries to the expansion phase were much earlier in German DOM (4,000–6,000 years ago) than in Korean MUS (2,000–4,000 years ago) and the difference may reflect the different history of agriculture between the two regions. In contrast, the population size of CAS was the smallest in ∼10,000–20,000 years ago but the reduction was much milder than that in the other subspecies. The milder population bottleneck would explain the higher genetic diversity of CAS.

This study clarifies the genome-wide relationship among *M. musculus* subspecies. Our *f*_3_ and *f*_4_ statistics support the close relationship between CAS and MUS, which is consistent with a study using dense genome-wide SNP data from wild-derived inbred mouse strains (White et al. 2009). In the presence of rampant gene flow between populations, inference of population splitting is extremely difficult. In particular, we found that the removal of obvious hybrid samples drastically changes the topology of evolutionary relationship of subspecies (Figure 5). Additional analyses using different types of data (*f*_3_ statistics and MSMC, Figure 4 and 6) supported that the CAS-MUS clade is most likely. We would like to make a caution that the bifurcating tree construction methods that do not assume any gene flow among taxon potentially give a biased result if hybrid samples are included in the analysis.

*M. musculus* has been widely used as an animal model of evolutionary genetic and biomedical research. Revealing the genetic background and evolutionary history of this species substantially contributes to the knowledge of these models among research communities. In this study, we report 98 novel whole-genome sequences of wild house mice, which have been collected from a range of regions focusing on the Eurasian continent and surrounding islands. Our analysis captured the genetic diversity of wild house mice on the Eurasian continent, which has not been well studied, and revealed a complex pattern of admixture among the three major subspecies. The high-quality whole-genome sequencing data presented in this study will be important for future studies in evolutionary ecology, population dynamics and natural selection by introgression among subspecies using wild house mice.

## Materials and Methods

### Materials

We collected 98 wild house mouse samples from a wide range of the Eurasian continent and remote islands. These samples are identical to those studied by Li et al. 2021, who reported and analyzed whole mitochondrial genome sequences. In addition, whole-genome sequencing data from 35 wild house mice and 8 western Mediterranean mice (*M. spretus*, SPR), reported by Harr et al. 2016, were downloaded from the European Nucleotide Archive (PRJEB9450, PRJEB11742, PRJEB14167, PRJEB2176, and PRJEB11897). In this study, the mitochondrial haplogroup of each sample was determined using the data of Li et al. 2021. The mitochondrial haplogroups in the Harr et al. data were determined by constructing a multiple sequence alignment with the data of Li et al. 2021 using Muscle implemented in Mega7. Supplementary Table S1–S3 lists the detailed information regarding the samples, and Supplementary Figure S1 illustrates their geographical sampling locations.

### Mapping genomic read pairs and single-nucleotide variant calling

For the 98 wild house mouse samples, paired-end sequences 100-bp in length were determined using the BGISEQ-500 platform (Li et al. 2021). The quality of reads was checked and visualized using FastQC (Andrews 2010) and MultiQC (Ewels et al. 2016) programs.

All raw reads were mapped to the GRCm38 (mm10) house mouse reference genome sequence using the bwa-mem algorithm with the “-M” option (Li and Durbin 2009). The Samblaster program with “-M” option was used for marking PCR duplicate reads (Faust and Hall 2014). For publicly available datasets, we used 43 samples with a median coverage >20. Raw SNV and insertions/deletion (indel) calls were performed using the GATK4 HaplotypeCaller program with the “-ERC GVCF” option (McKenna et al. 2010). All genomic variant call format (gVCF) files were merged using the GenomicDBImport function, and the variants of all samples were jointly called using the GenotypeGVCFs function.

Raw SNVs and indels were processed using the GATK4 Variant Quality Score Recalibration (VQSR). VQSR is a machine learning process that uses known variants as a training dataset and predicts whether a new variant would be a true positive or a false positive. In order to run GATK4 VQSR, we used the “mgp.v3.snps.rsIDdbSNPv137.vcf.gz” and “mgp.v3.indels.rsIDdbSNPv137.vcf.gz” files downloaded from the webserver of the Sanger Institute (ftp://ftp-mouse.sanger.ac.uk/REL-1303-SNPs_Indels-GRCm38/) as training datasets for SNVs and indels, respectively. We also included HARD-filtered SNV data as a training dataset. A HARD filtering process for SNVs was performed using the following parameters: QD < 2.0, FS > 60.0, MQ < 40.0, MQRankSum < −12.5, and ReadPosRankSum < −8.0. Moreover, a HARD filtering process for indels was performed using the following parameters: QD < 2.0, FS > 200.0, InbreedingCoeff < -0.8, ReadPosRankSum < -20.0, and SOR > 10.0. We assumed that SNVs and indels within the 90% tranche (90% acceptance in all reliable training SNVs datasets) were true positive SNVs and indels, and we used them for downstream analyses. VQSR-passed SNVs were further filtered according to mappability to the *Mus musculus* reference genome, computed using the GenMap software (Pockrandt et al. 2020). We computed mappability scores using the “-K 30” and “-E 2” options and analyzed sites with mappability values of 1. Mappability filtering retains highly unique regions in the reference genome.

Because we could not reliably distinguish the sexes of some of our samples from morphological records, we assigned the sexes of samples based on the read depth coverages on the X and Y chromosomes (Table S2). We used Samtools Depth to calculate the coverage of each sample in the non-pseudoautosomal regions of the sex chromosomes that passed the mappability filter. The ratios of average X-chromosomal to Y-chromosomal coverages showed a clear bimodal distribution, 0.96–1.15 and 7.43–195.20, where samples within the former and latter ranges likely represent male and female samples, respectively.

Kinship inference among samples was performed using KING software with the “—kinship” option (Manichaikul et al. 2010). According to KING software, the expected ranges of kinship coefficients are >0.354 for duplicate/monozygotic twins, >0.177 and <0.354 for first-degree relationships, >0.0884 and <0.177 for second-degree relationships, >0.0442 and <0.0884 for third-degree relationships, and <0.0442 for unrelated individuals. The 13 samples (12 *Mus musculus* samples and 1 *Mus spretus* sample) were excluded with relationships closer than third-degree. In total, 128 samples (94 of our samples and 34 public samples) were kept after filtering process.

Synonymous and nonsynonymous variants were assigned using the SnpEff (Cingolani et al. 2012b) and SnpSift (Cingolani et al. 2012a) programs with house mouse gene annotation data version “GRCm38.101 (ftp://ftp.ensembl.org/pub/release-101/gtf/mus_musculus/)”. This calculation was done by counting the number of synonymous and non-synonymous variants on gene-by-gene basis.

### Population structure analysis

Using VCFtools (Danecek et al. 2011), SNVs were further filtered and converted to PLINK format (Purcell et al. 2007) containing only biallelic autosomal SNVs, retaining sites successfully genotyped in all samples. Typically, SNVs in linkage disequilibrium are excluded from analysis of population structure; however, we did not eliminate these SNVs because our mouse samples were highly structured at the subspecies level and eliminating these SNVs would have left too few SNVs for analysis. Principal component analysis (PCA) was performed using the smartpca program (Patterson et al. 2006). Default parameter settings were used with the exception of declining to remove outlier samples. The color codes in Figure 1a and 1b were assigned according to Maxwell’s color triangle. Outgroup *f*_3_ statistics were computed using Admixtools, employing the SPR population as an outgroup with the “outgroupmode” option (Patterson et al. 2012). The *f*_4_ statistics were also computed using Admixtools with the “f4 mode” and “printsd” options (Patterson et al. 2012). ADMIXTURE (Alexander et al. 2009) software was used for population stratification. We computed cross-validation error values (--cv option) from *K* = 1 to *K* = 5 for datasets that either included or excluded SPR. The identity-by-state (IBS) distance matrix between all pairs of individuals was calculated using the PLINK “--distance 1-ibs” option, and the matrix was used to construct a neighbor-joining tree (Saitou and Nei 1987) using the Ape package in R (Paradis et al. 2004).

### Demographic inference using PSMC and MSMC/MSMC2

Pairwise sequentially Markovian coalescent (PSMC) analysis was performed for the sampled individuals (Li and Durbin 2011). To obtain a consensus autosomal genome sequence for each individual, which is a required input for PSMC, the “mpileup” samtools command was applied to the dataset using “-C 50, -O, -D 60, -d 10” options. The PSMC analysis options (-t and -p) were chosen according to the default settings suggested by the PSMC software. The time interval parameters were set to “4+25*2+4+6” with 25 iterations. To validate for the variance of population size *N*_e_, we performed 100 replications of bootstrap method for each subspecies representative samples.

The MSMC (Schiffels and Durbin 2014) version 2 software (MSMC2: https://github.com/stschiff/msmc2) (Schiffels and Wang 2020; Malaspinas et al. 2016) was used to estimate the effective population size (*N*_e_) changes and subspecies divergence time. In our MSMC/MSMC2 analysis, we performed estimations using phased haplotype sequence as input. We estimated phased haplotype by ShapeIt4 software (Delaneau et al. 2019). We used the data “Mapping Data for G2F1 Based Coordinates” from “Mouse Map Comverter (http://cgd.jax.org/mousemapconverter/)” to provide the recombination rate input file for MSMC2. Mappability was taken into account and non-unique sequence positions were not used for calculations. The time interval parameters were set to “1*2+50*1+1*2+1*3” with 20 iterations. To estimate the subspecies divergence time, we used two haplotypes from each population totaling four haplotypes to calculate rCCR for the input of MSMC2. According to Shiffels et al. 2014, the rCCR variable ranges between 0 and 1 (in some cases, the calculation is unavoidably greater than 1), and the value close to 1 indicates that two populations were one population at the point of time. Heuristically, the half value of rCCR, i.e. rCCR = 0.5, is estimated as the timing of the separation of the two populations. The bootstrapping of MSMC2 was done by cutting the original input data into blocks of 5 Mb each, and then randomly sampling them to artificially create a 3-Gbp-length genome. Calculations were performed on a total of 20 of these artificially created data sets. In order to estimate *N*_e_ over time, eight haplotypes from four samples of each MUS (KOR01–03 and 05) and DOM (DEU01, DEU03, DEU04, and DEU06) population, and four haplotypes from two samples of CAS (IND03, IND04) were used for calculation. For estimation of subspecies divergence, we used total four haplotypes from two populations for each combination of CAS-MUS (IND04 and KAZ01), DOM-MUS (DEU07 and KAZ01), and CAS-DOM (IND04 and DEU07).

## Data Availability

The short-read sequence data generated in this study have been submitted to the DDBJ BioProject database (https://www.ddbj.nig.ac.jp/bioproject/) under accession number PRJDB11027. The datasets, parameter setting files, and scripts required for reproducing the analysis are deposited to Dryad digital repository (https://datadryad.org/) under doi:10.5061/dryad.66t1g1k1j.

## Acknowledgments

This work was supported by MEXT KAKENHI (grant 18H05511 to N.O.), Cooperative Research of the Inter-University Research Institute Corporation. This research was supported by Global Station for Big Data and Cybersecurity, a project of Global Institution for Collaborative Research and Education at Hokkaido University.

## Notes

### Competing Interest Statement

The authors have declared no competing interest.

### Summary of Updates

Revised Discussion about PSMC results.

