## Supplementary Figure for "Insights into *Mus musculus* population structure across Eurasia revealed by whole-genome analysis"

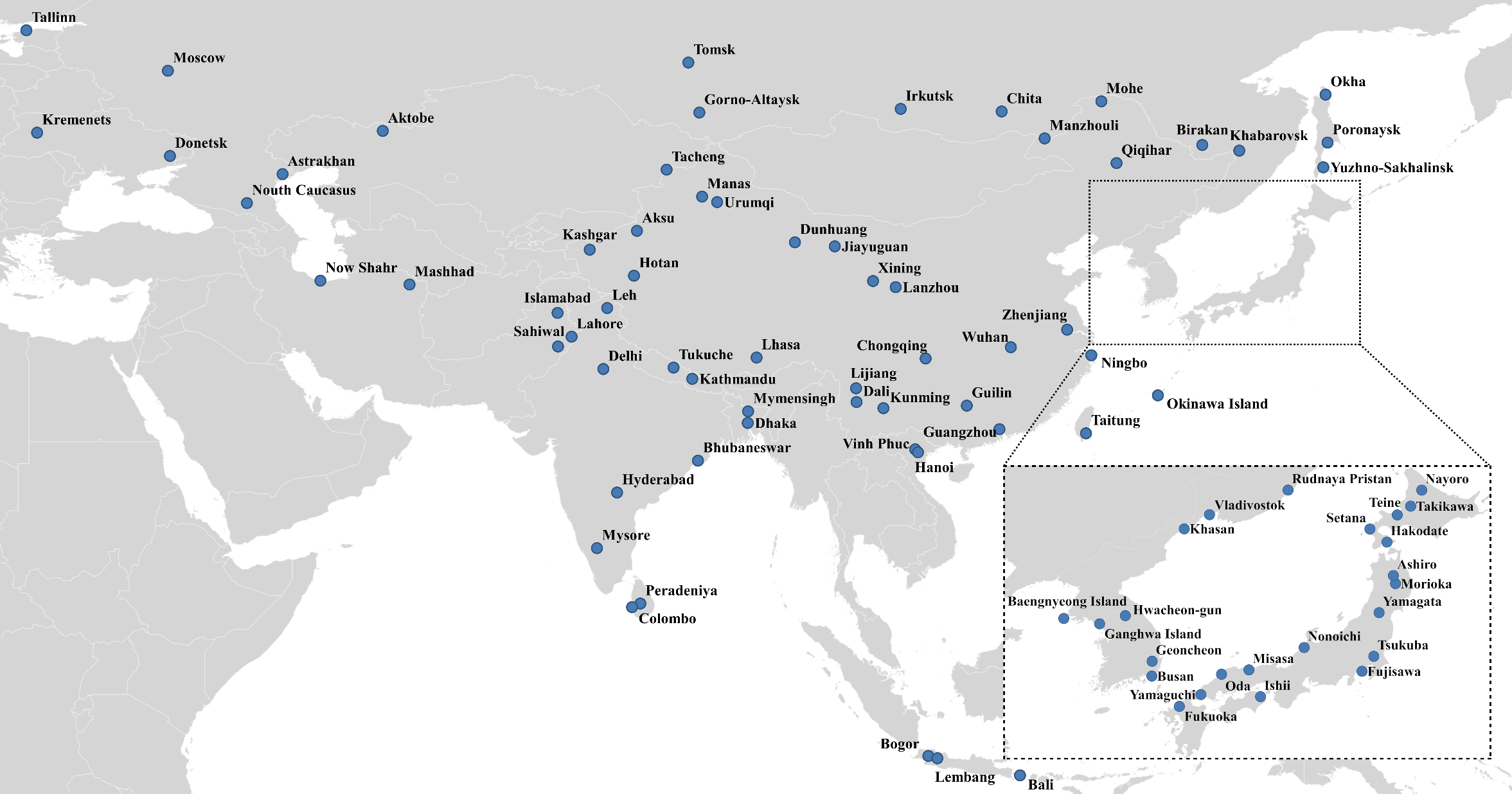

**Supplementary Figure 1**

Collection sites for wild house mouse samples used in this study.

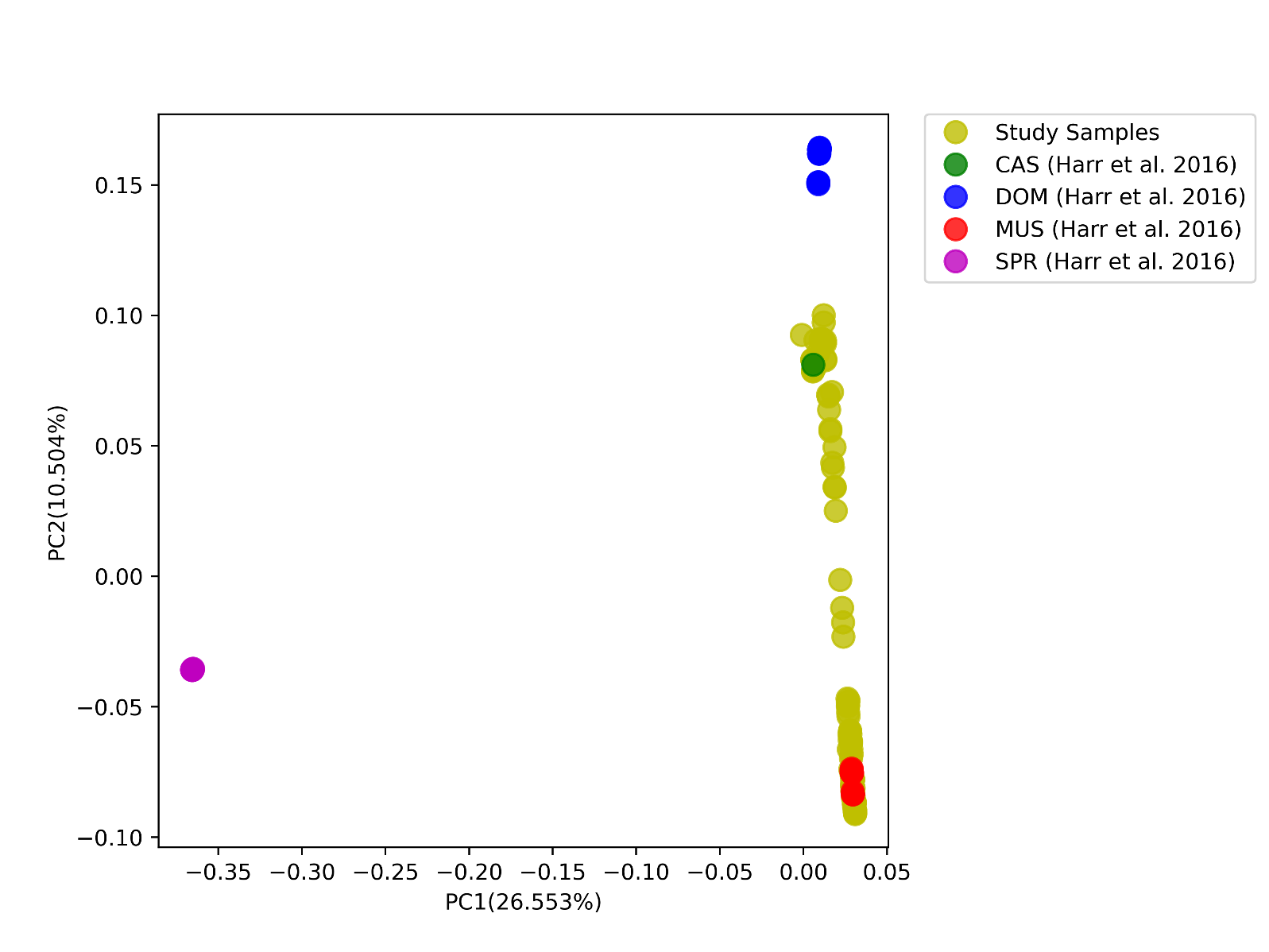

**Supplementary Figure 2**

The principal component analysis plot of house mouse *Mus musculus* and *Mus spretus.*  For comparison, the public data for MUS, CAS, and DOM samples are marked separately from our study samples.

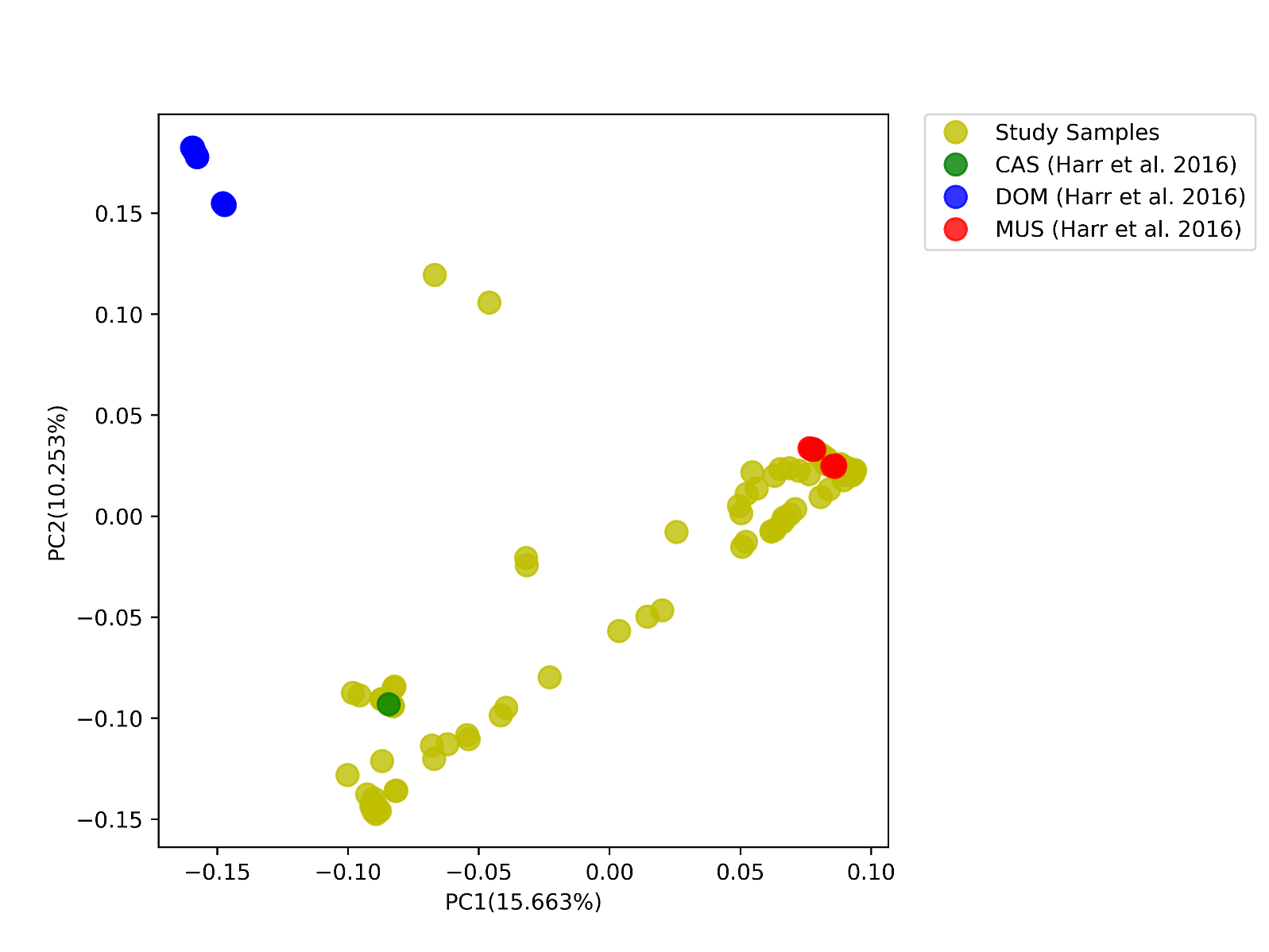

**Supplementary Figure 3**

The principal component analysis plot of house mouse *Mus musculus* used in this study. For comparison, the public data for MUS, CAS, and DOM samples are marked separately from our study samples.

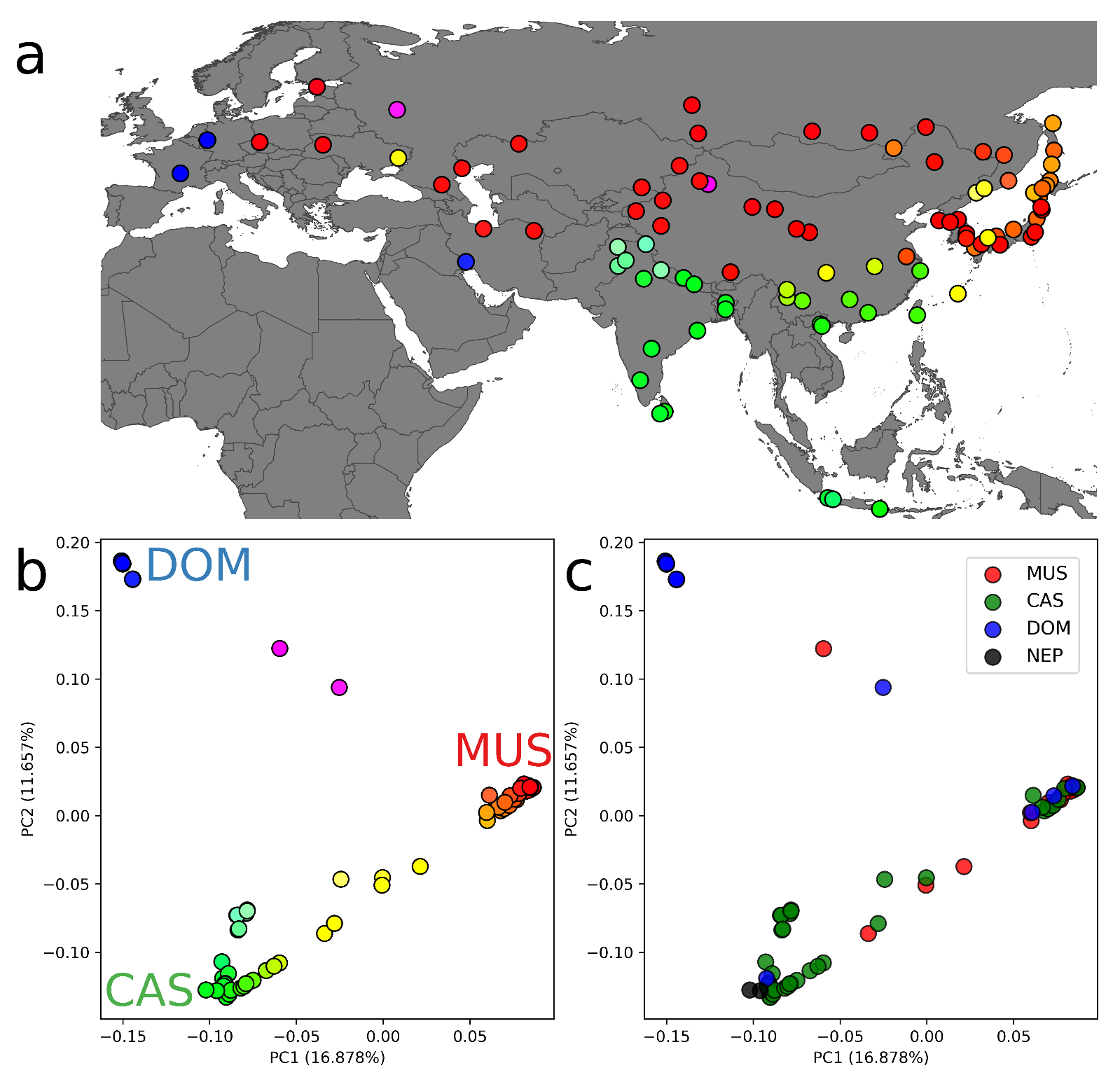

**Supplementary Figure 4**

The results of principal component analysis on the X chromosome in the wild house mice. A) A geographical map of sampling locations. The colors of circles correspond to the eigenvalues in panel B. The detailed names of sample collection sites are shown in Supplementary Figure 1. B) The PCA plot of the wild house mouse using X chromosome SNVs. The *x* and *y* axes represent the eigenvalues of PC1 and PC2, respectively. The circles were colored according to the “Maxwell’s Color Triangle” color scheme, assigning three vertices to the RGB colors. The red, green, and blue color intensities correspond to the MUS, CAS, and DOM genetic components, respectively. The proportion of variance for each eigenvalue is shown in parentheses on the axis label. C) The PCA plot of the wild house mouse using X chromosome SNVs, labeled with the mitochondrial genome haplogroup of each sample. The proportion of variance for each eigenvalue is shown in parentheses on the axis label.

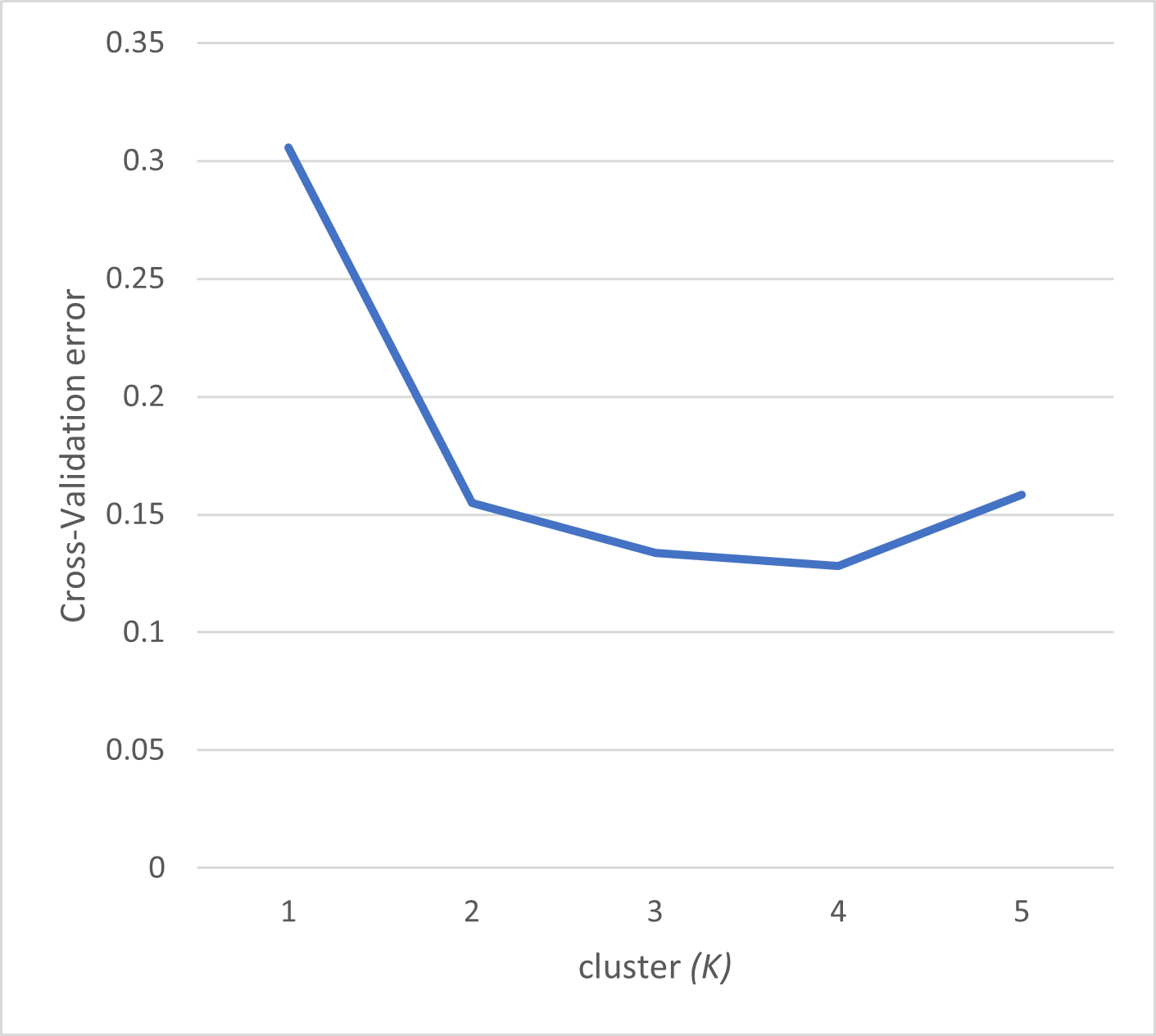

**Supplementary Figure 5**

The cross-validation error values of ADMIXTURE analysis. The better value of cluster *K* exhibits the lower cross-validation error value.

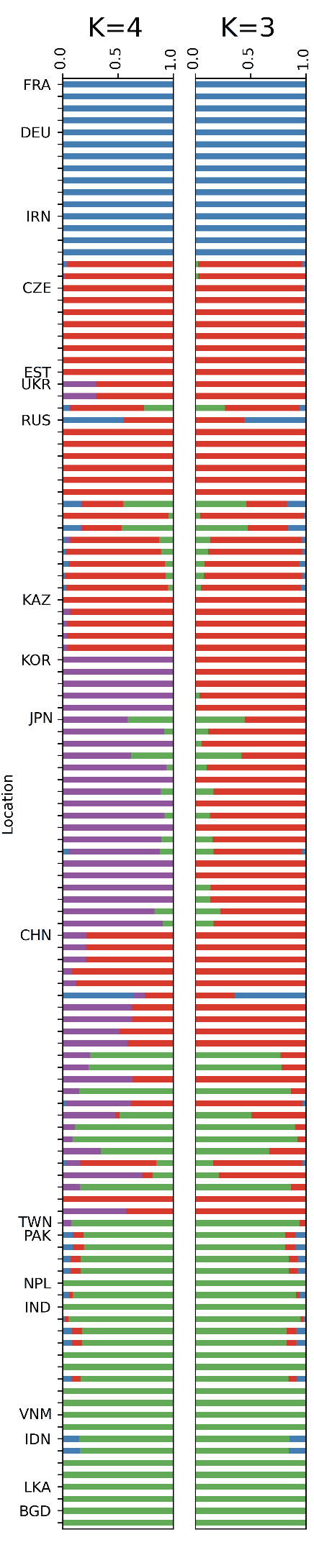

**Supplementary Figure 6**

The ADMIXTURE plot using X chromosomal data. The proportion of estimated subspecies genetic components are shown. The results of *K* = 3 and *K* = 4 are presented. In each country code, the samples are lined up according to longitude and latitude. From top to bottom, it represents samples from west to east and north to south.

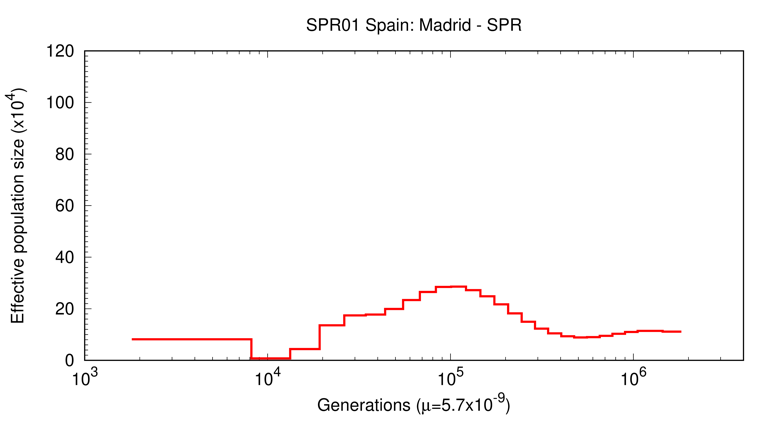

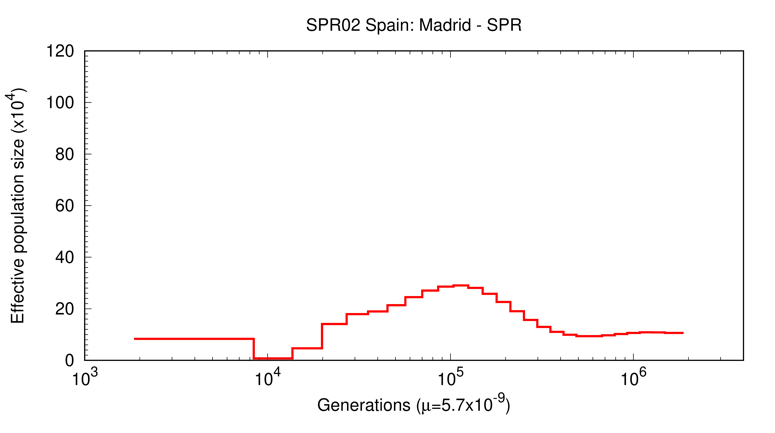

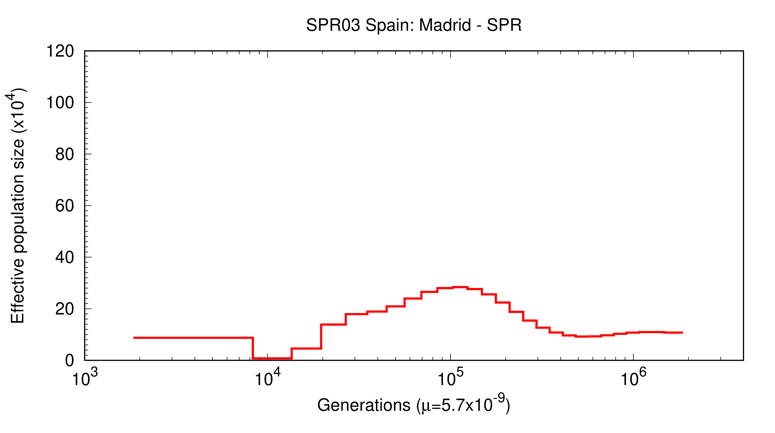

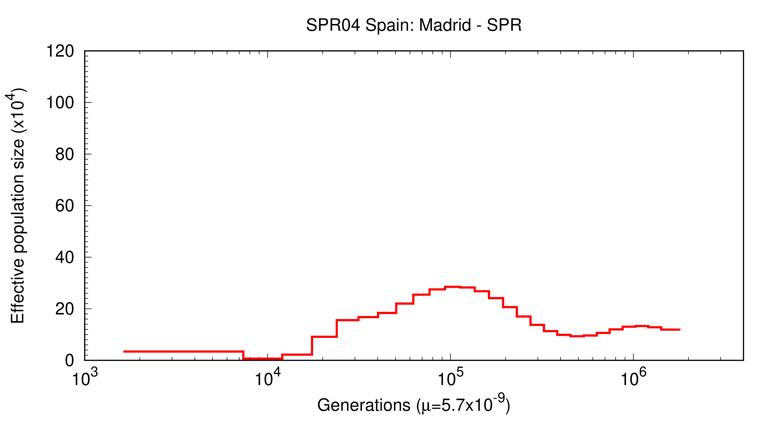

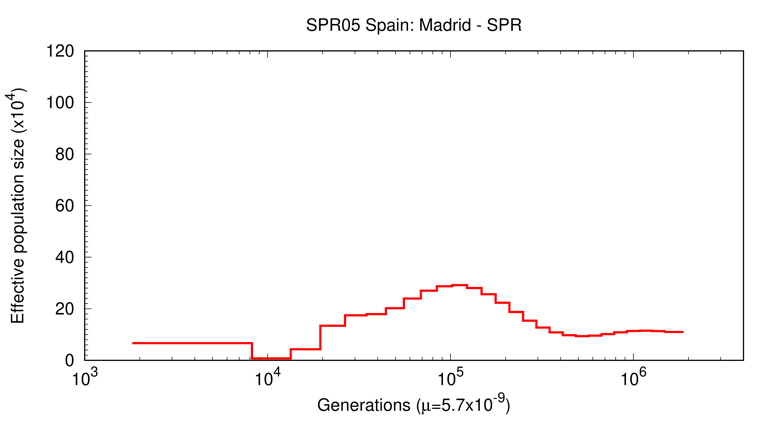

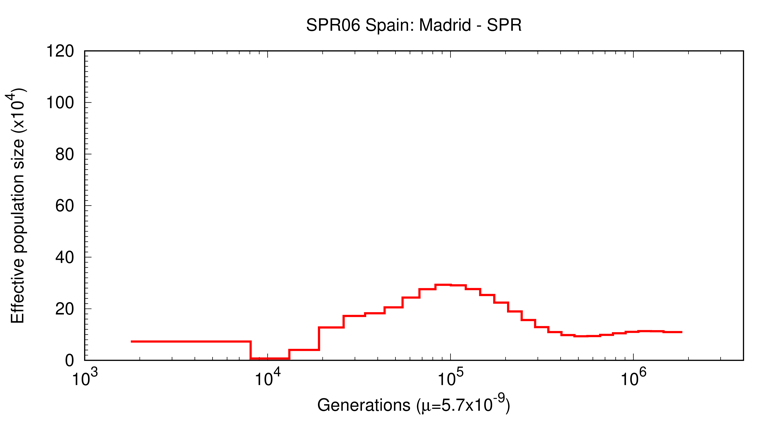

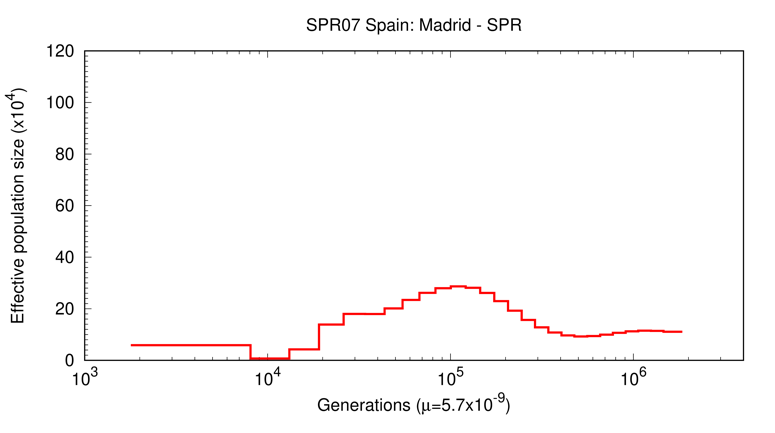

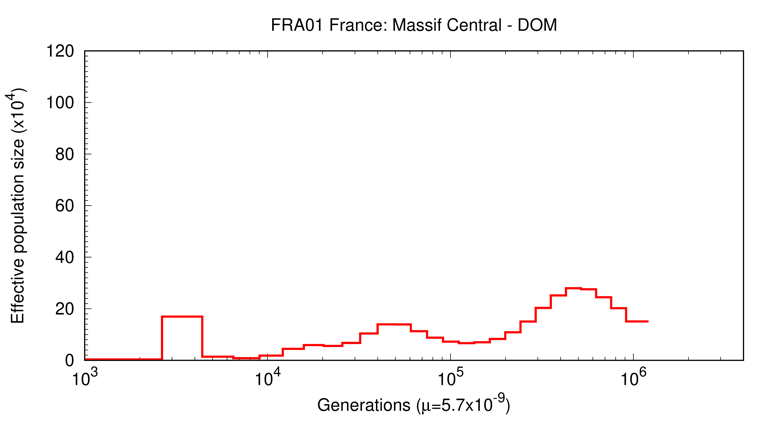

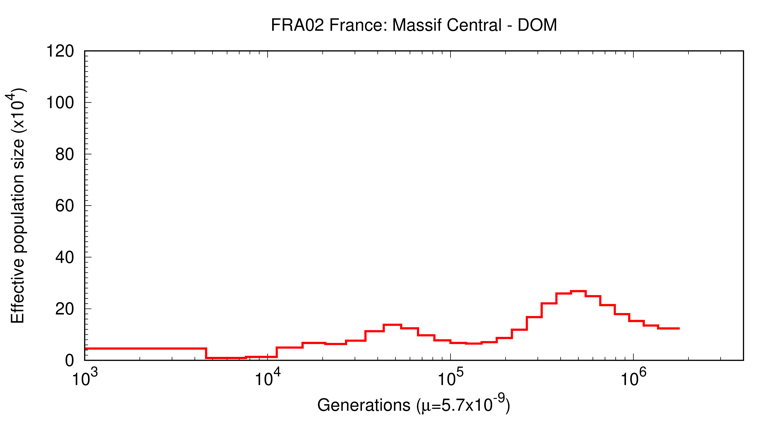

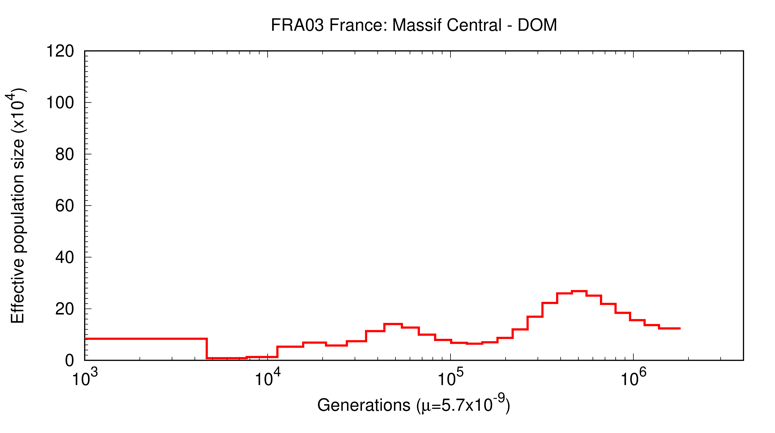

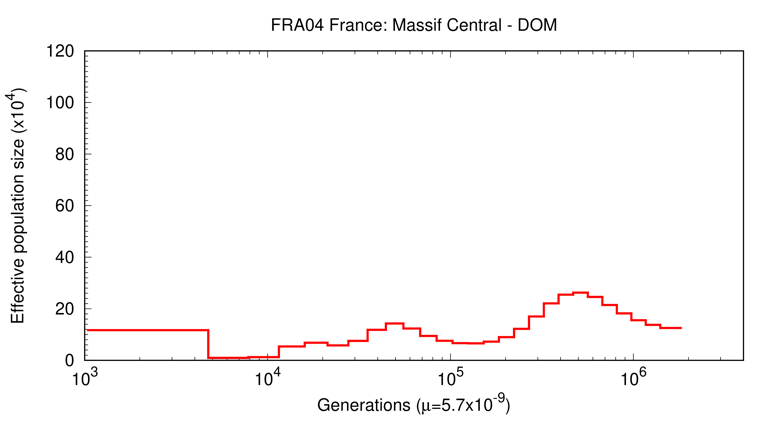

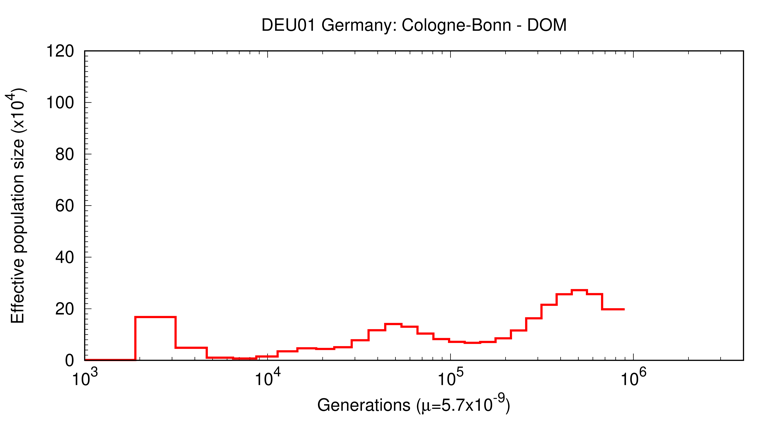

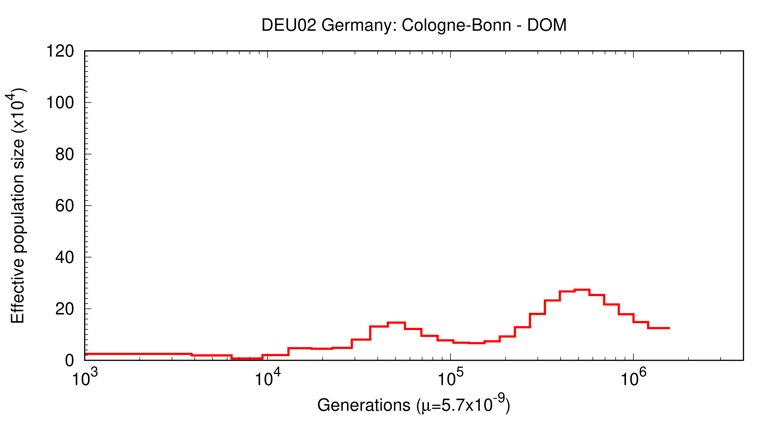

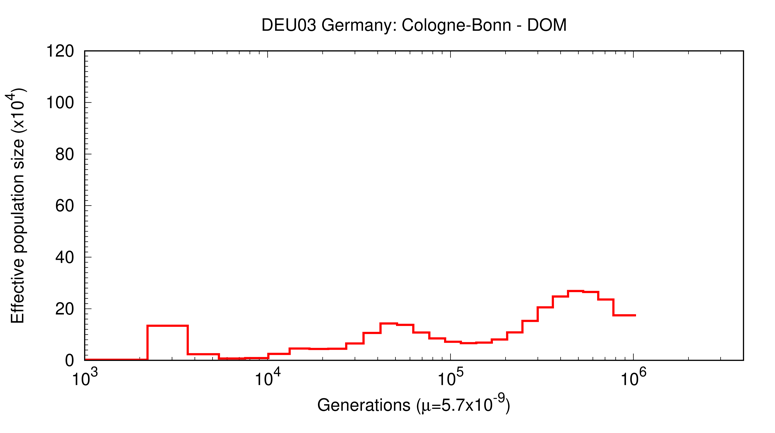

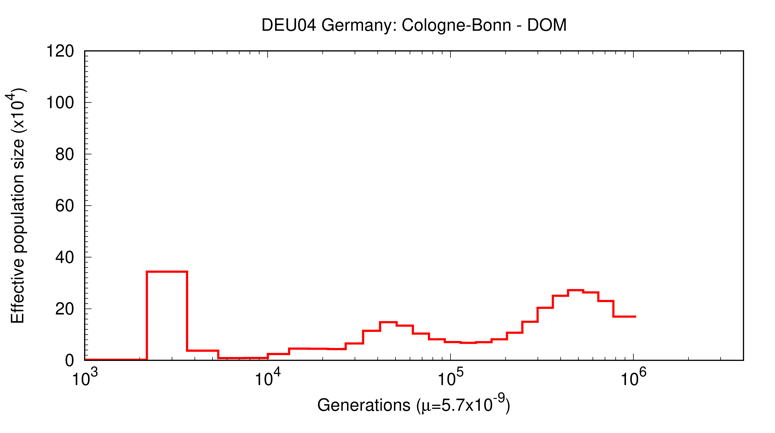

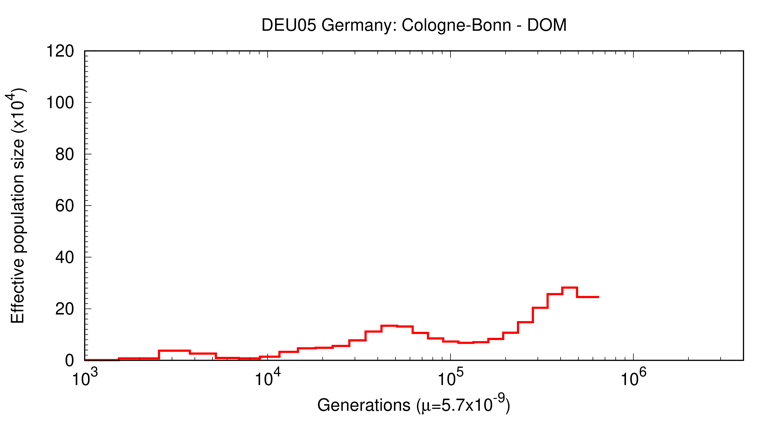

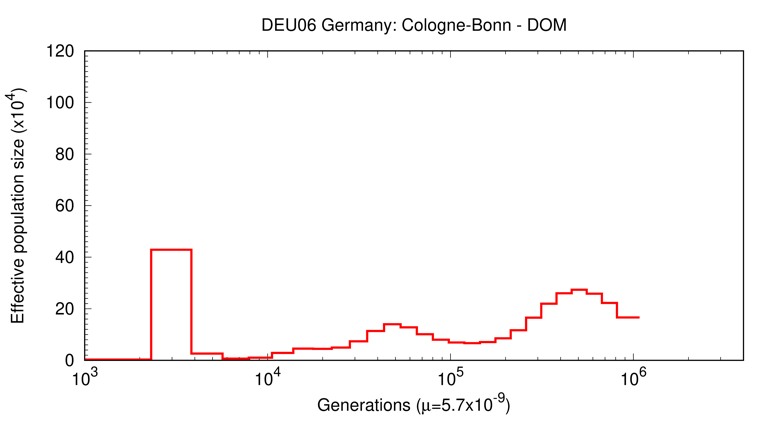

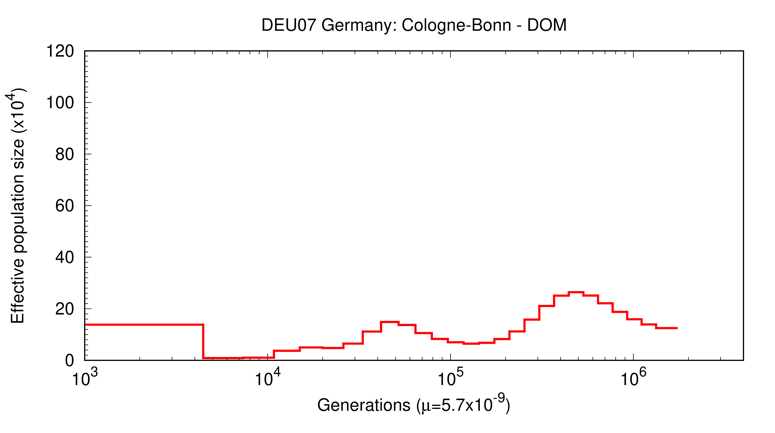

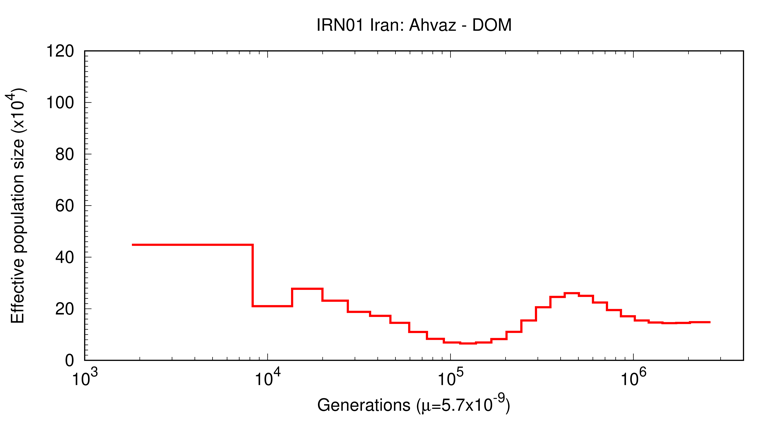

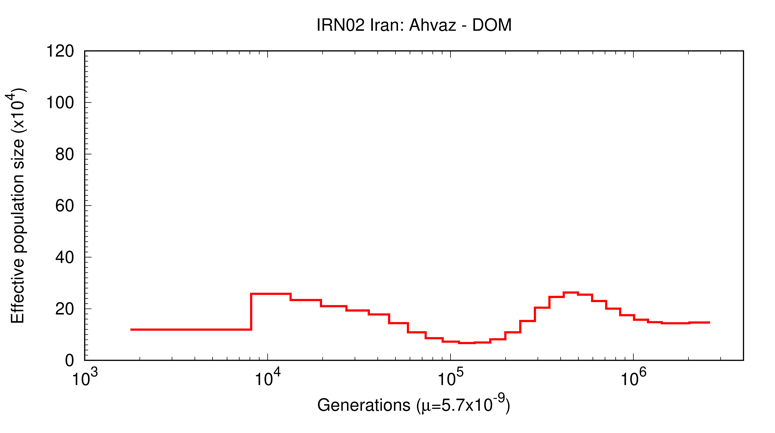

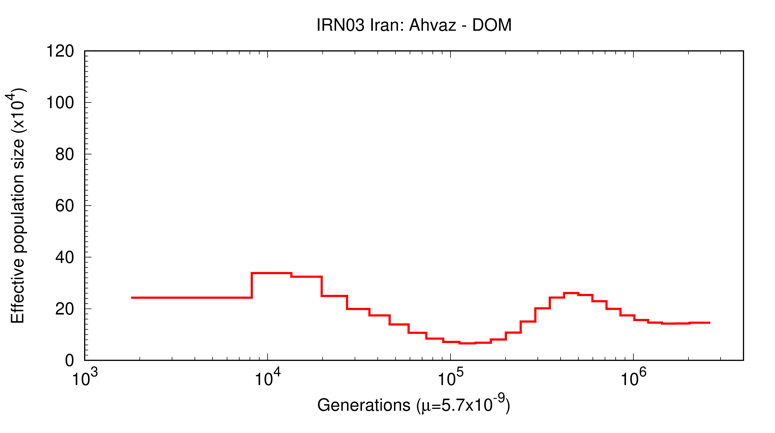

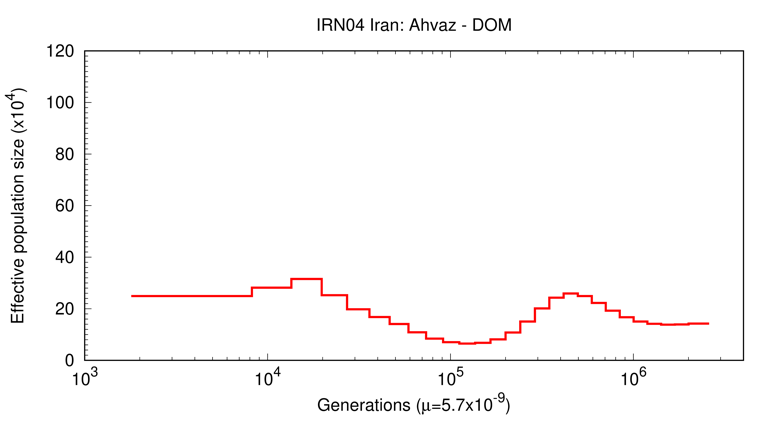

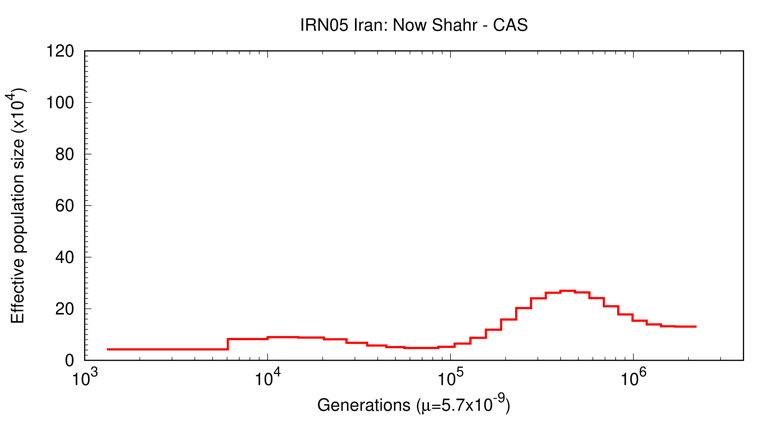

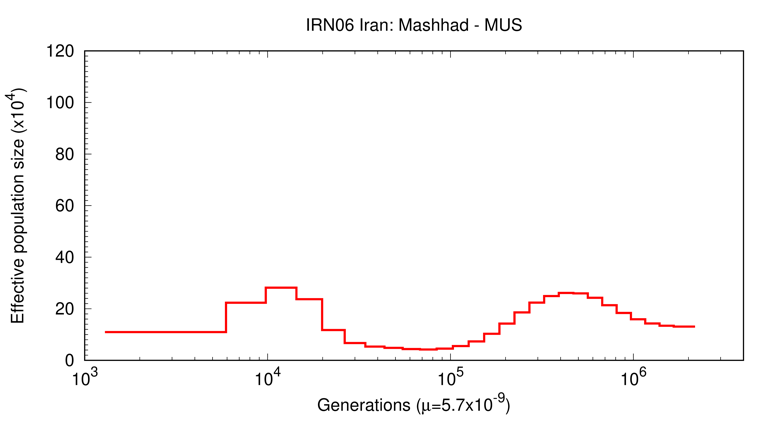

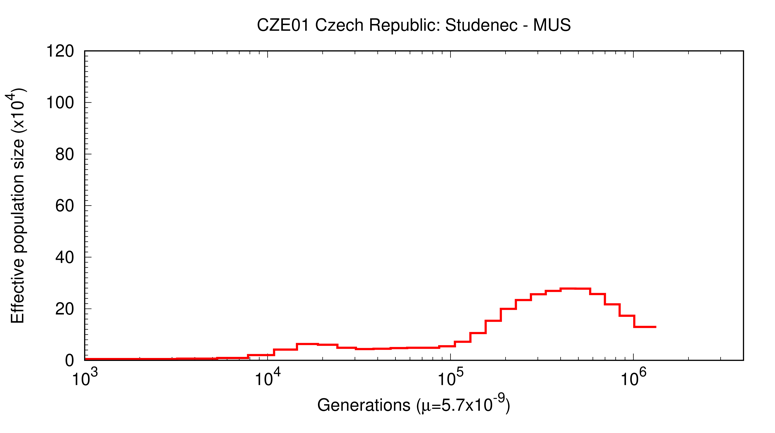

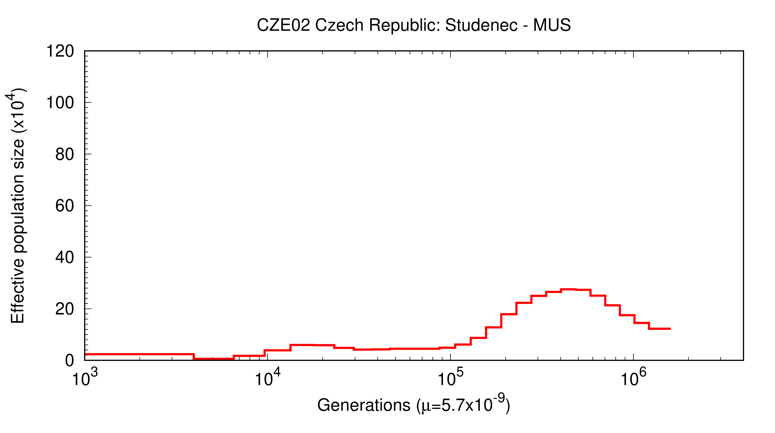

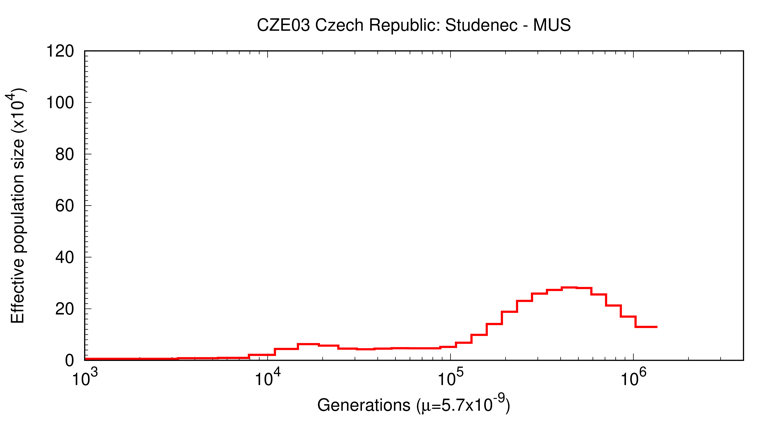

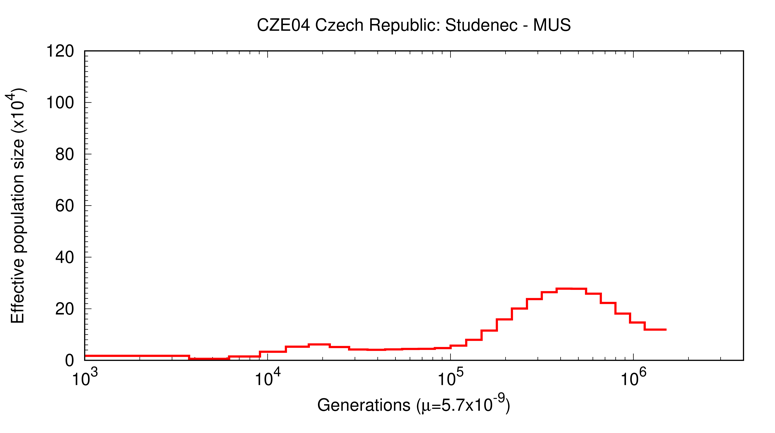

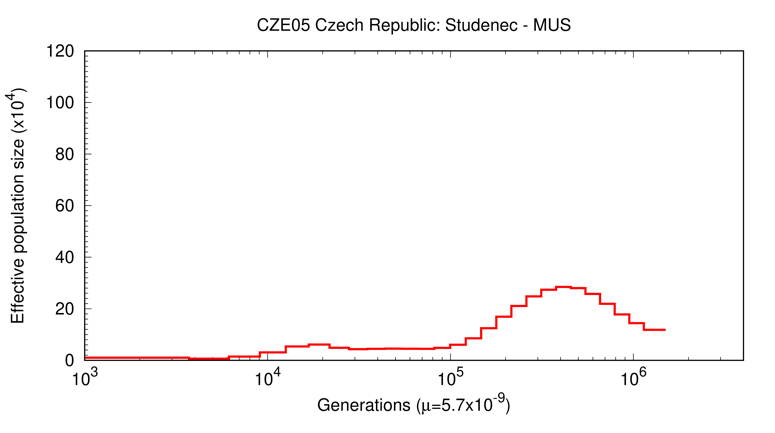

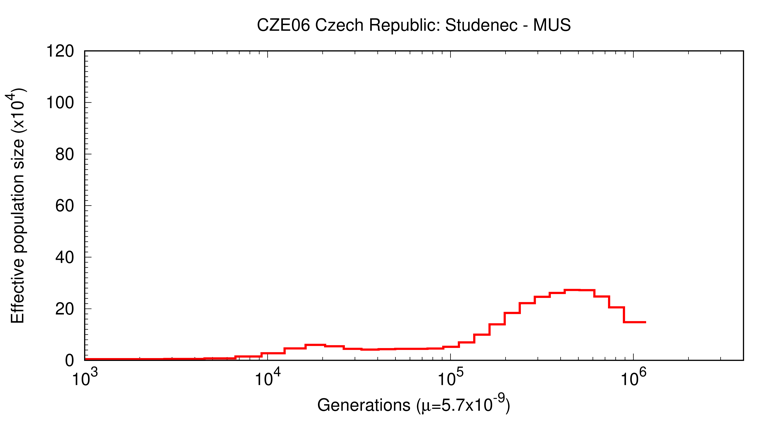

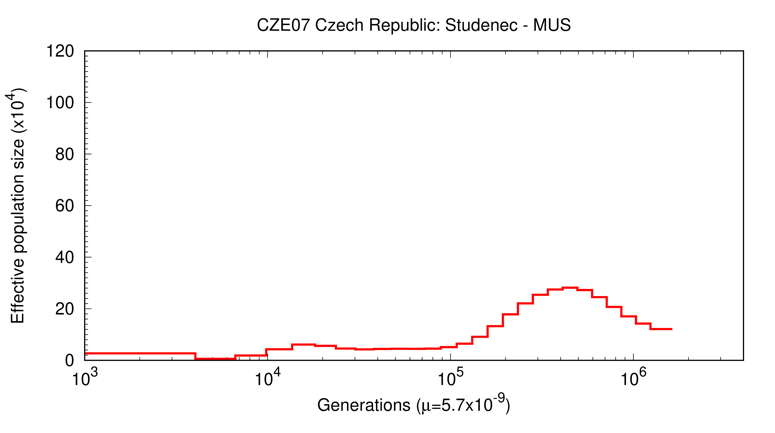

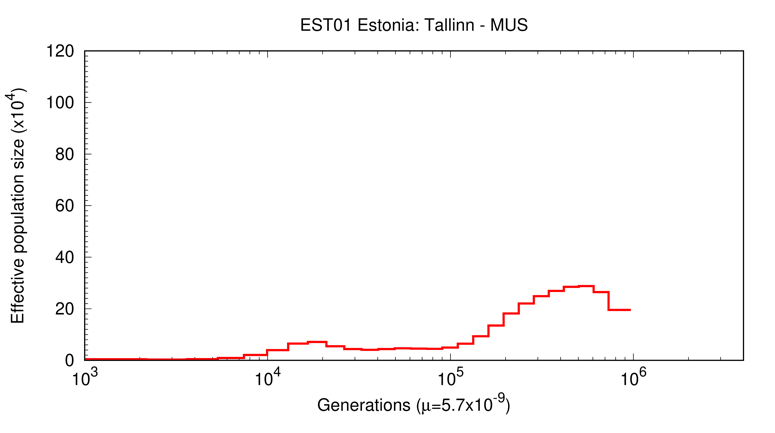

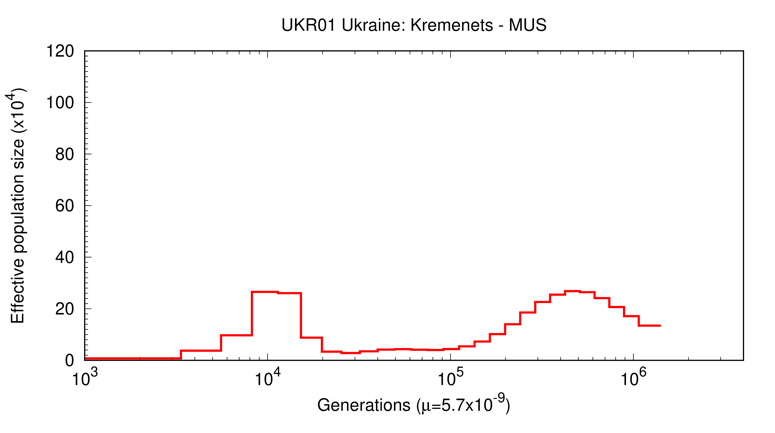

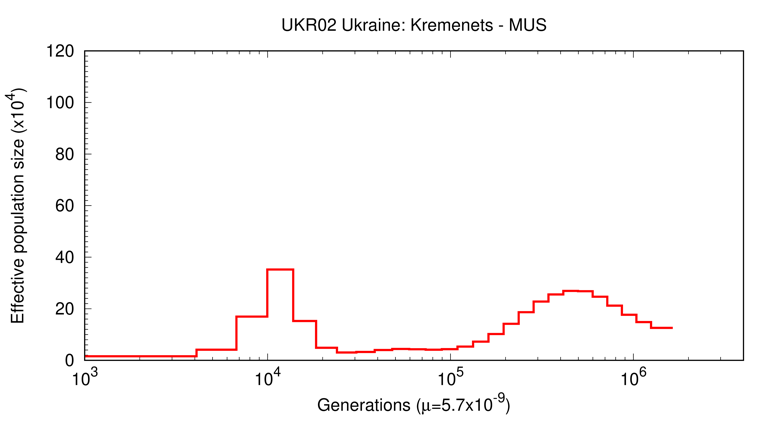

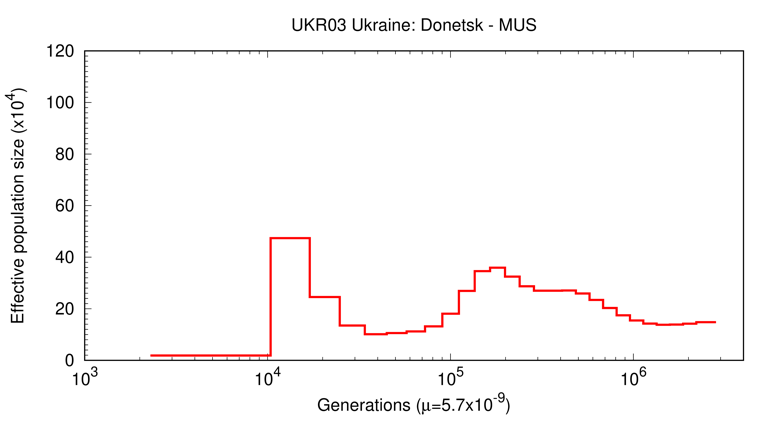

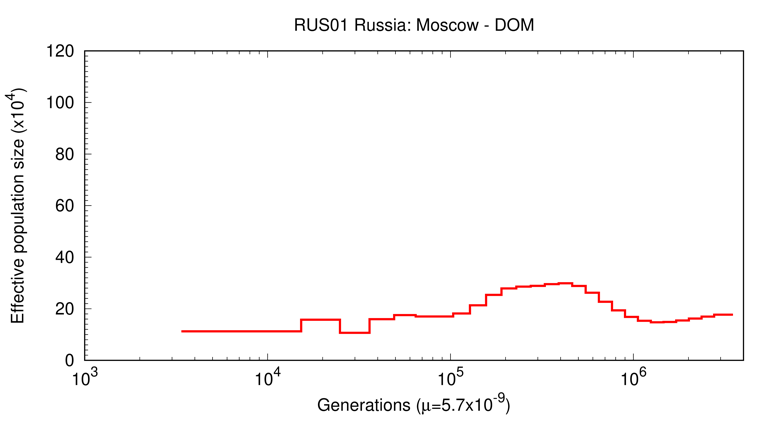

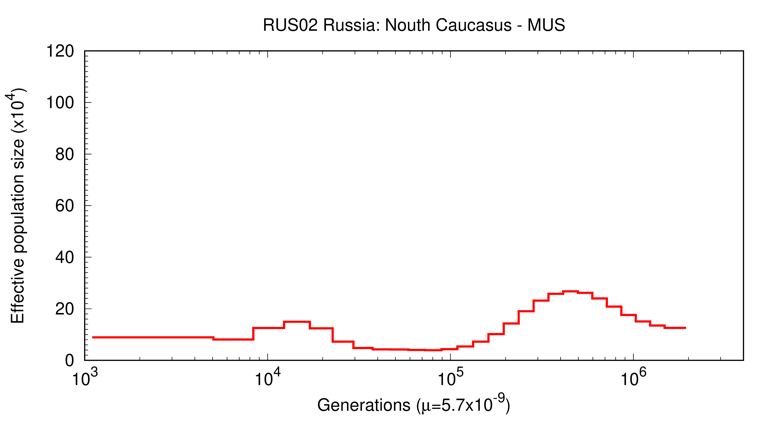

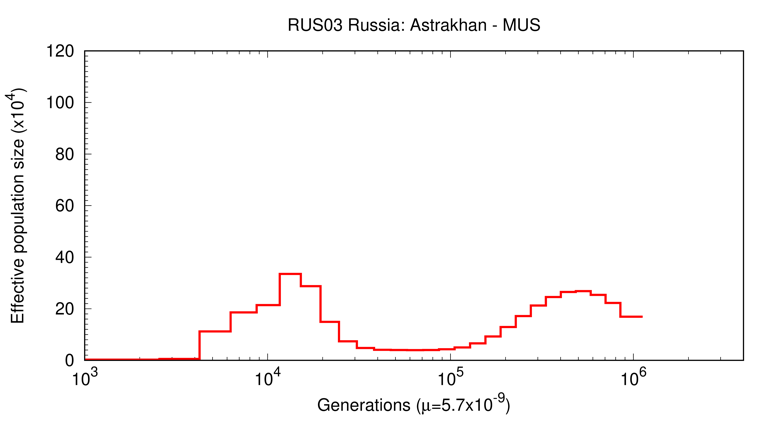

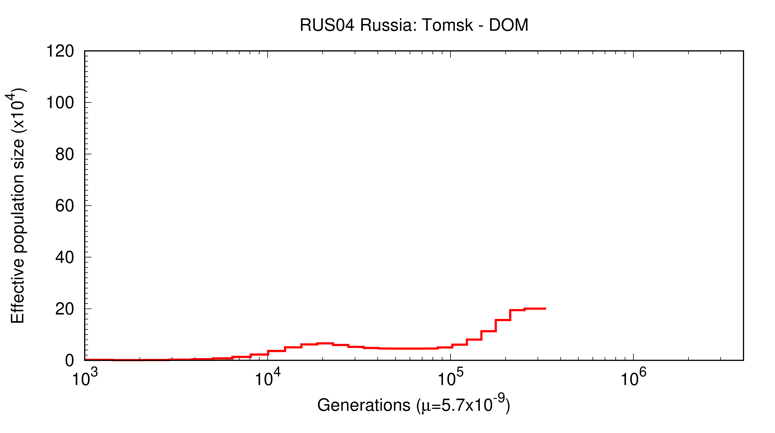

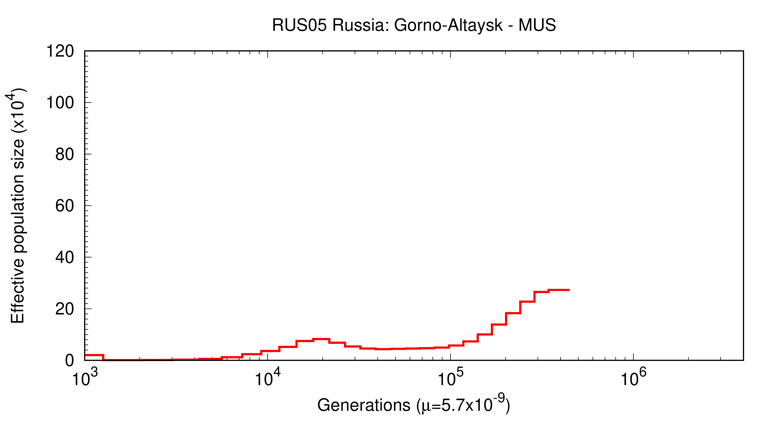

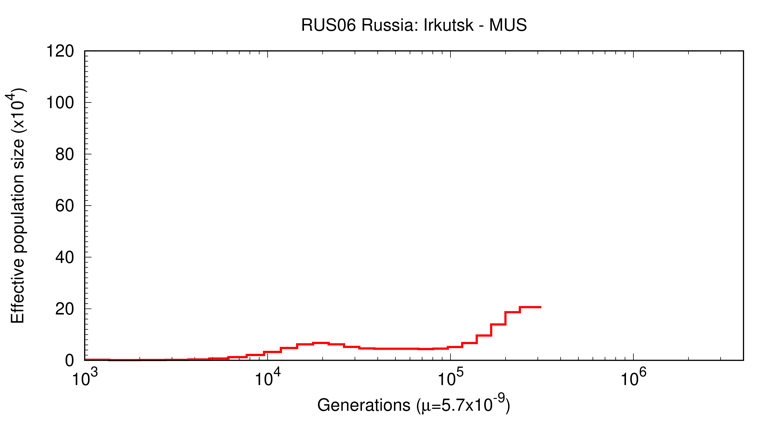

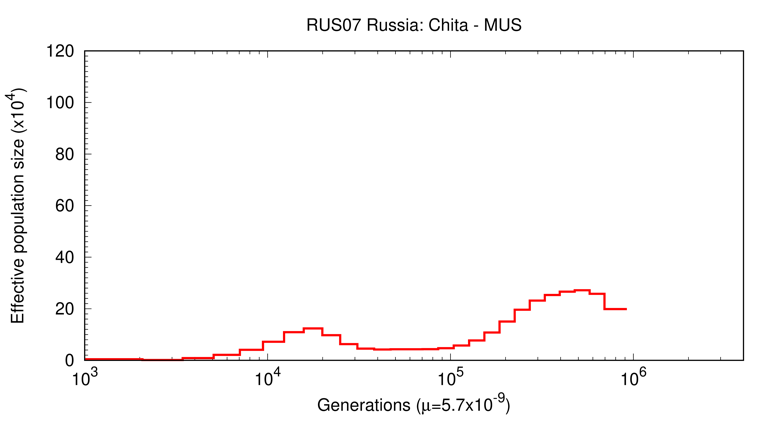

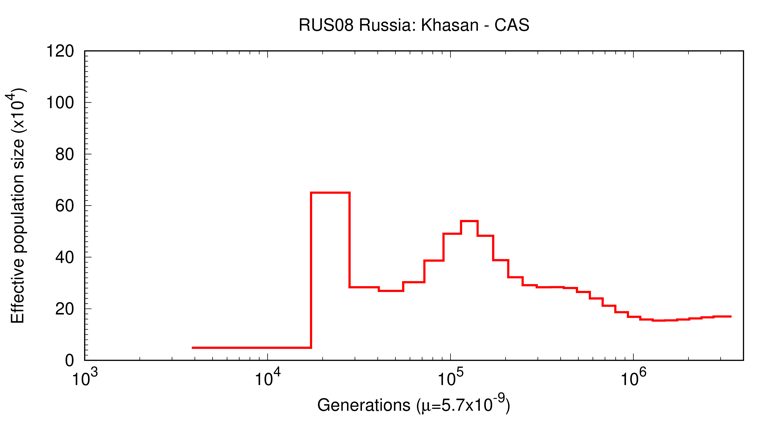

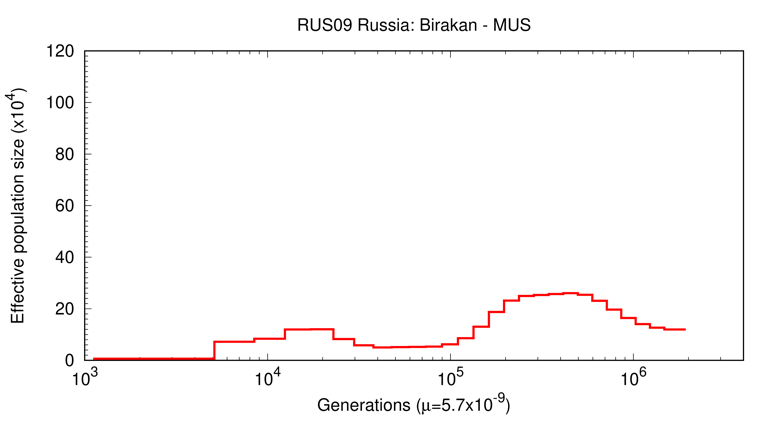

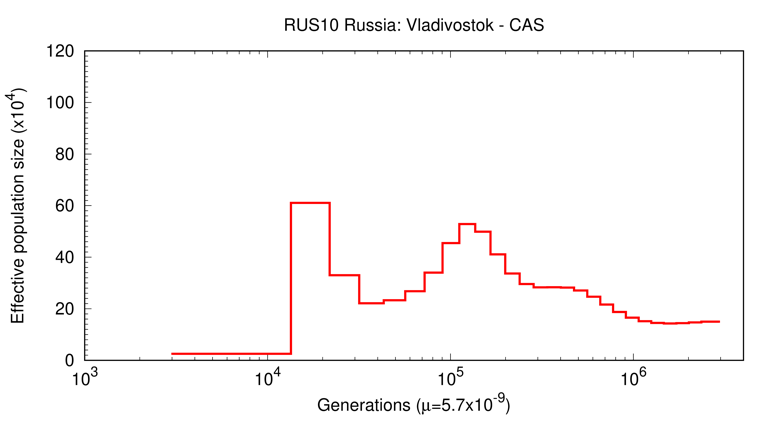

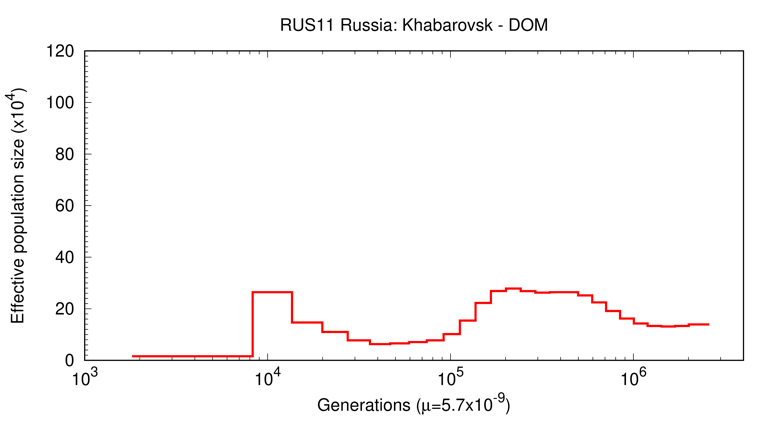

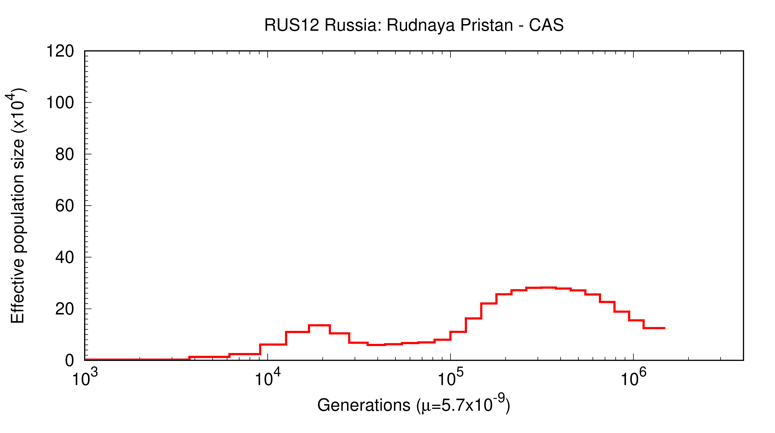

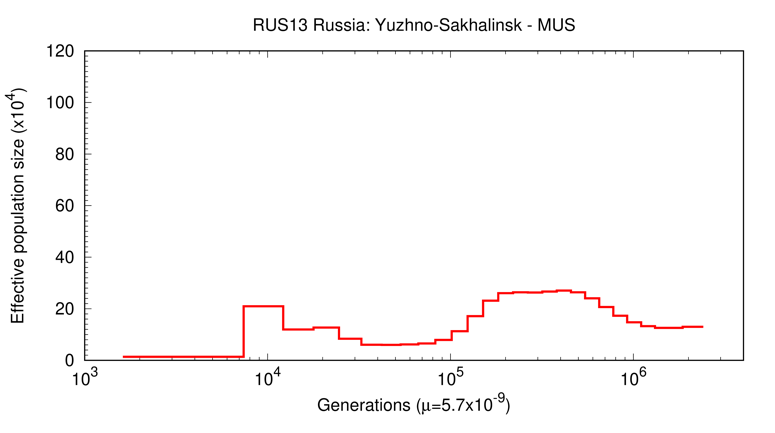

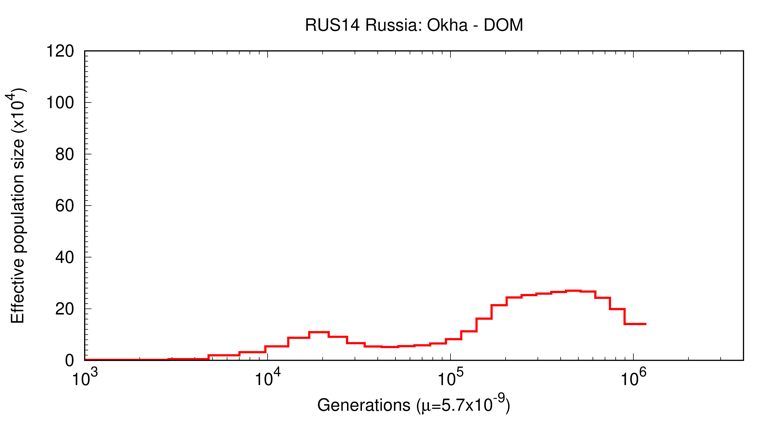

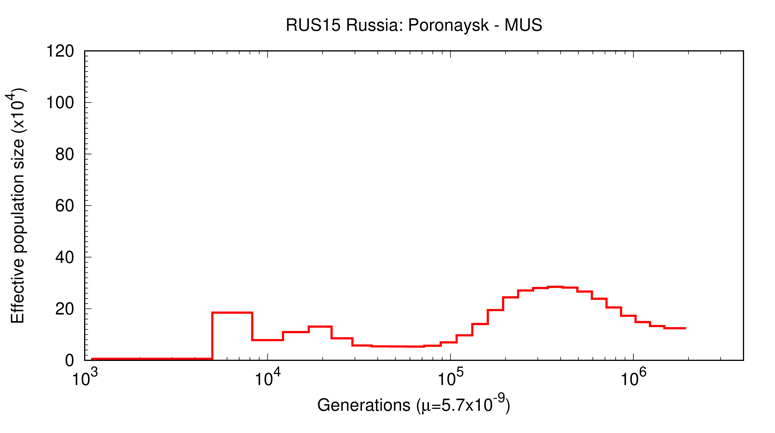

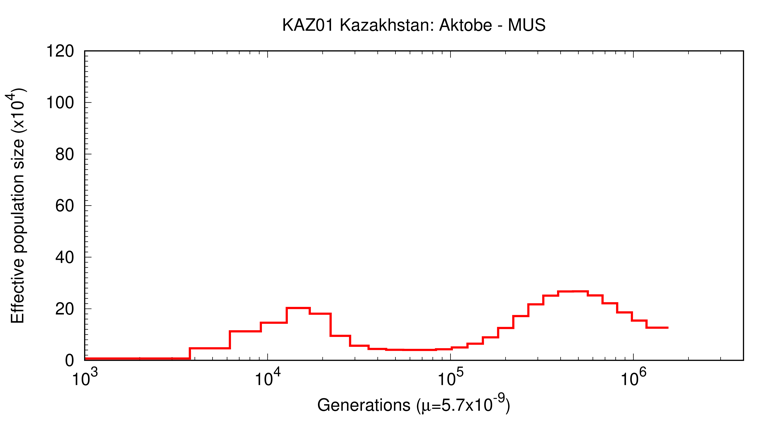

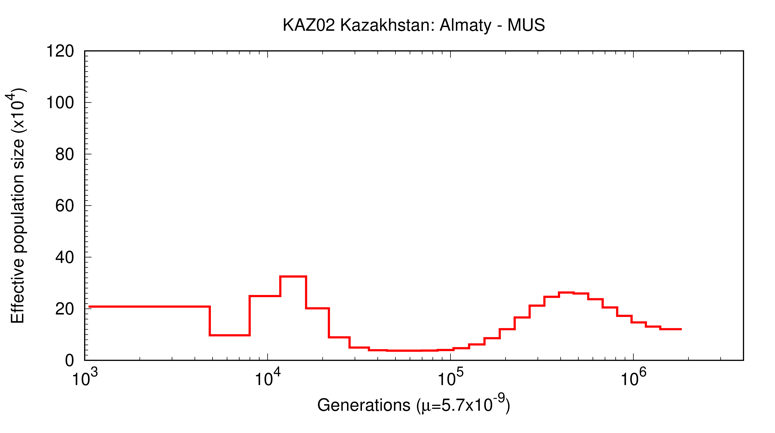

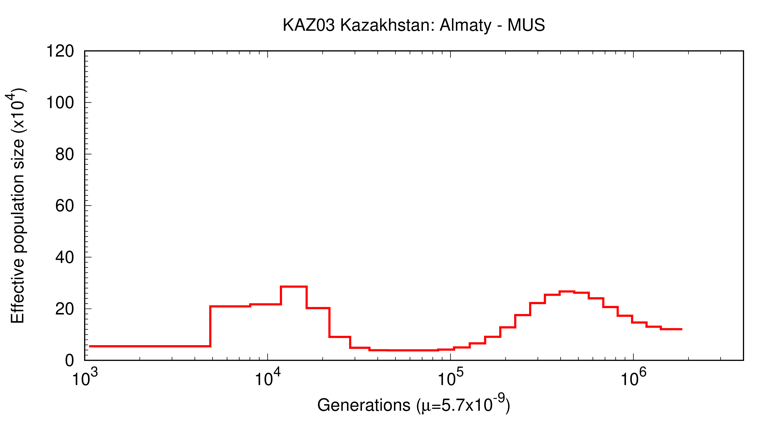

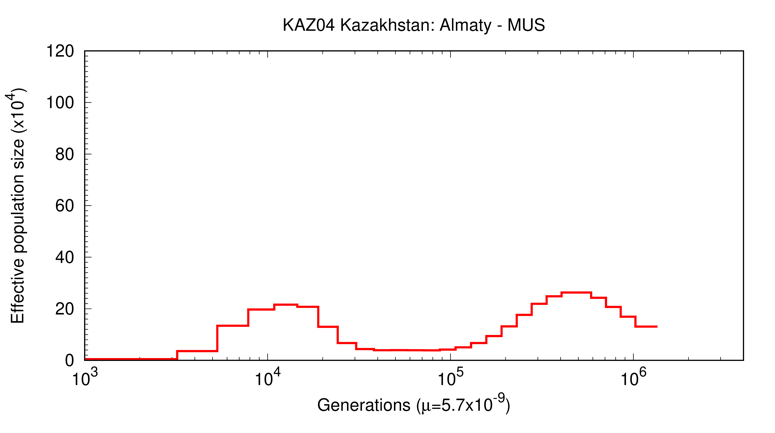

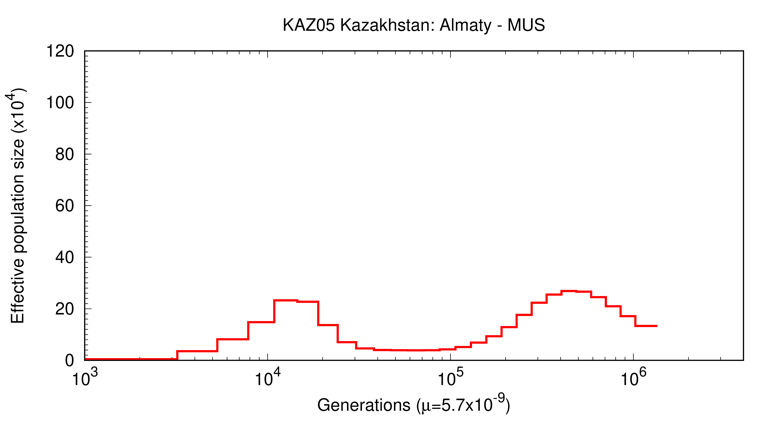

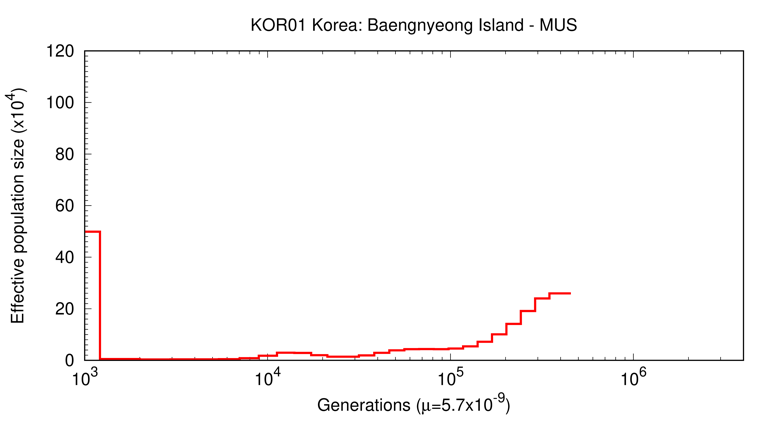

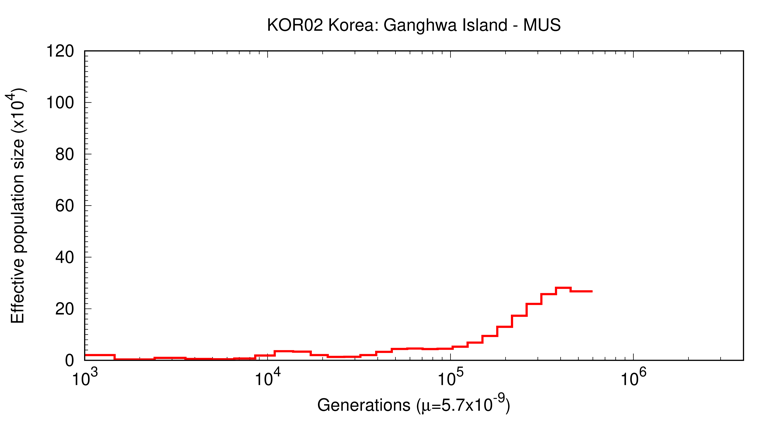

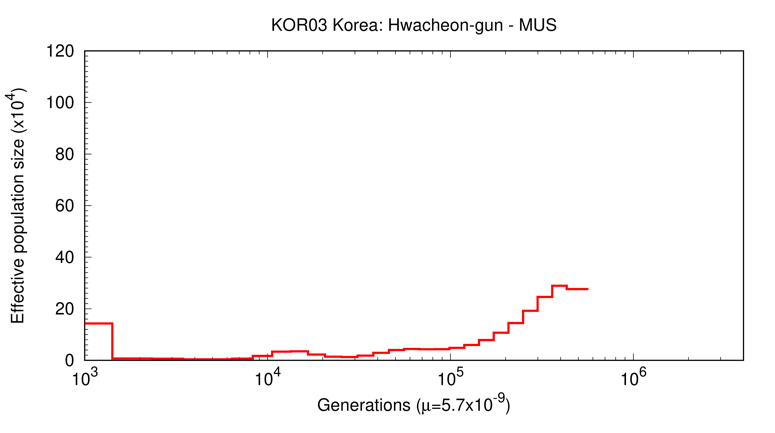

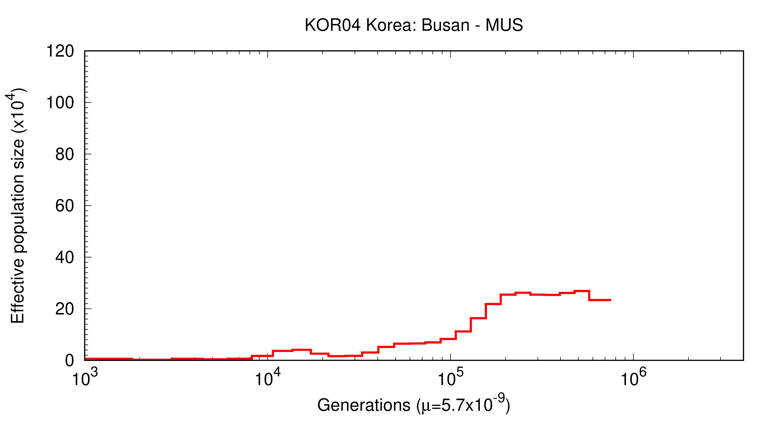

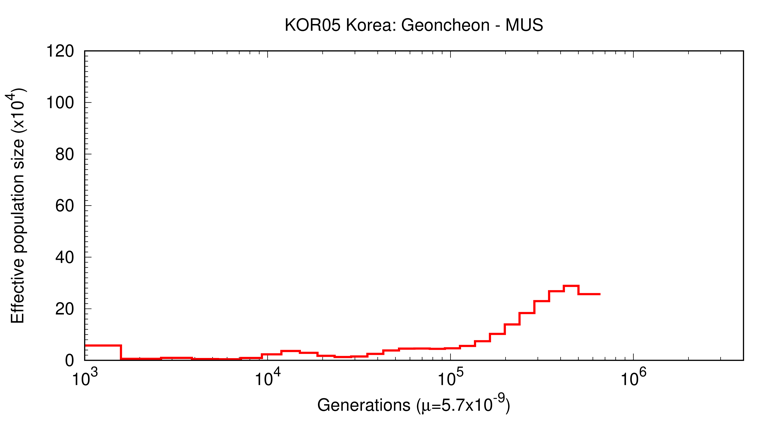

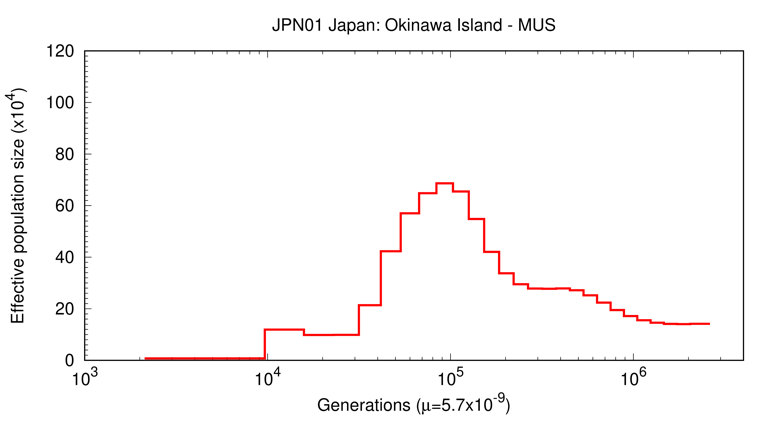

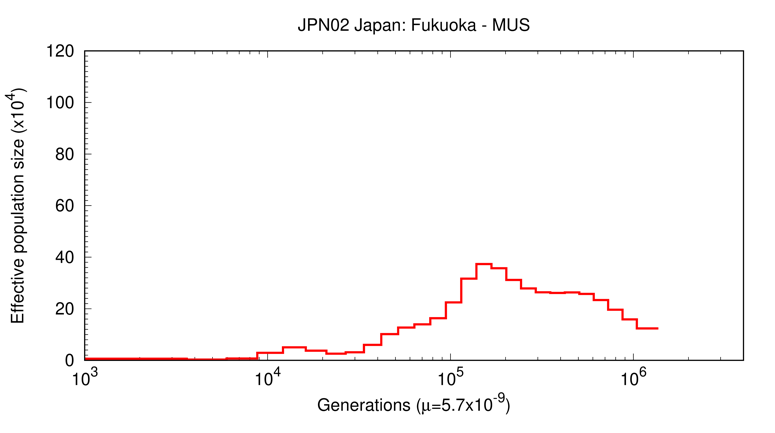

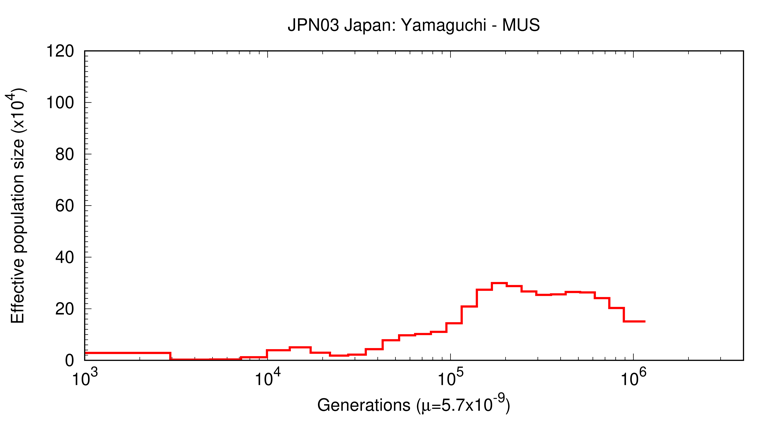

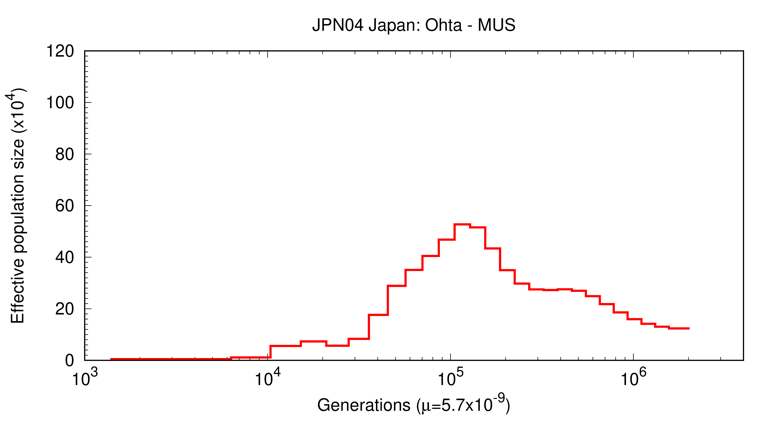

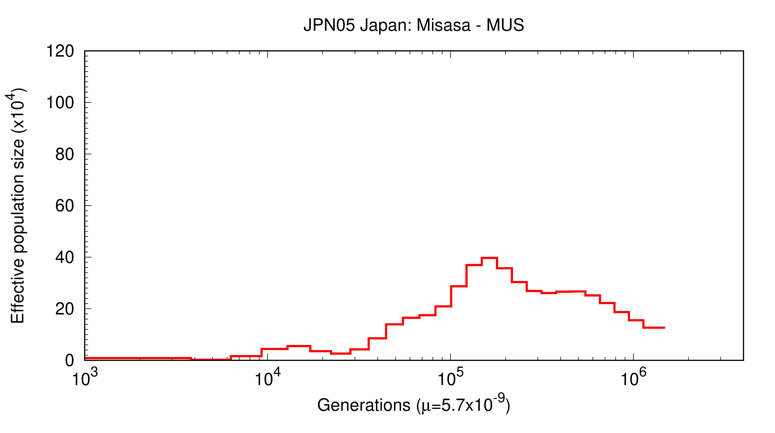

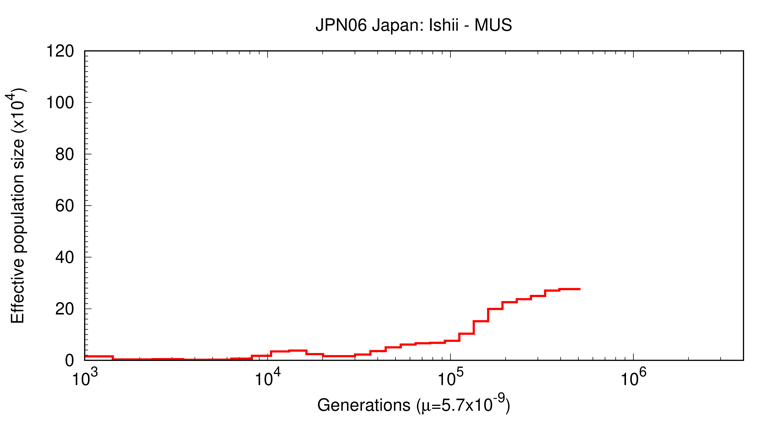

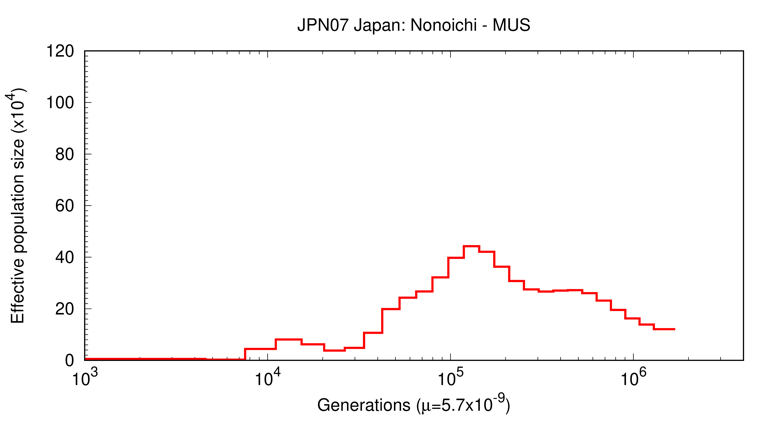

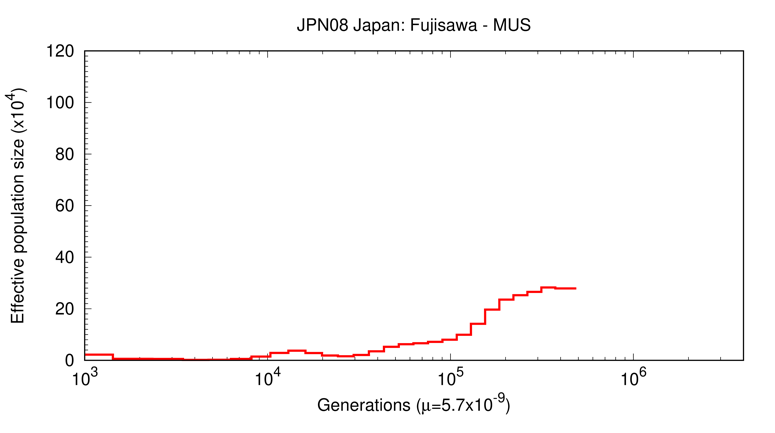

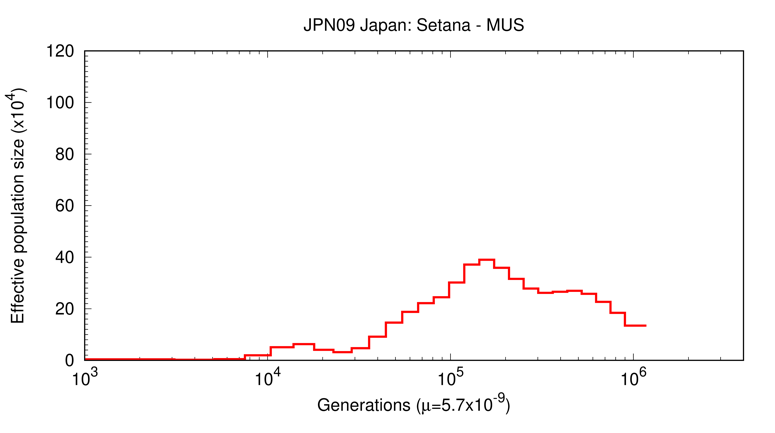

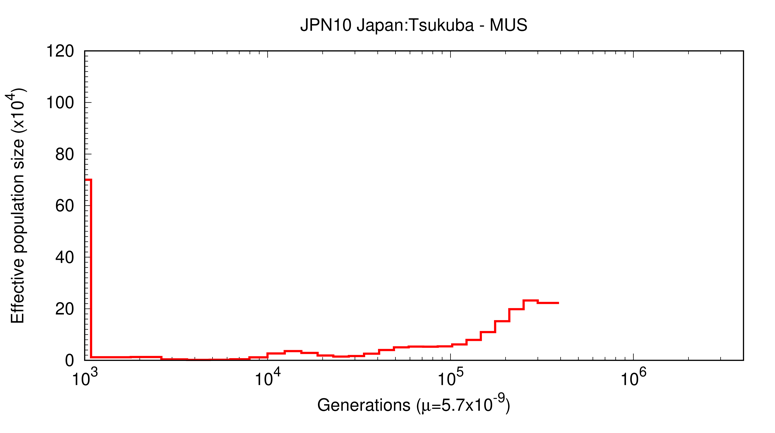

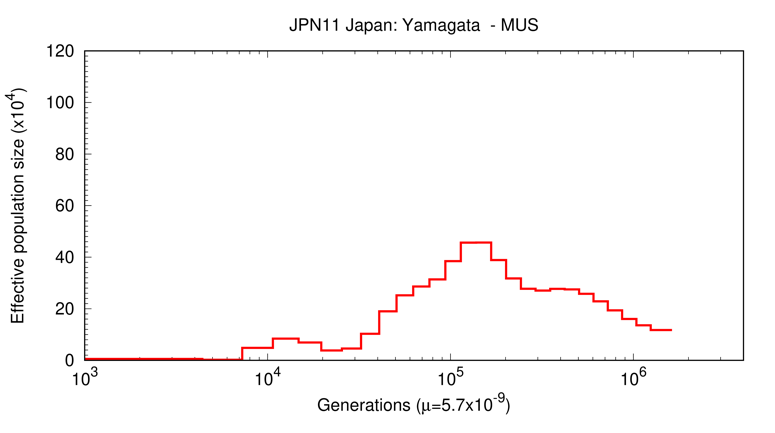

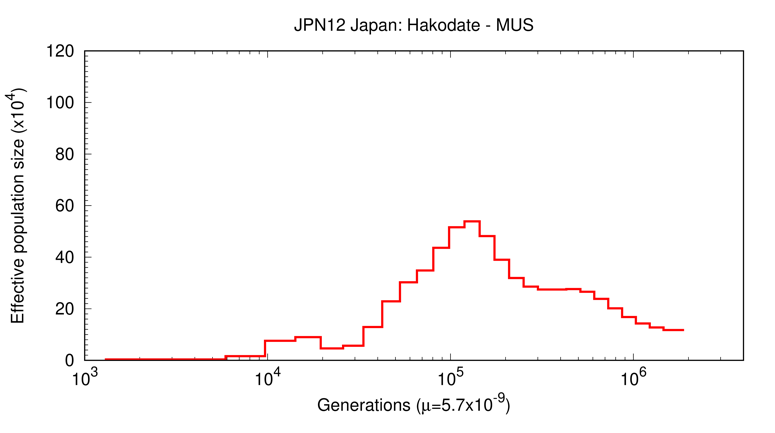

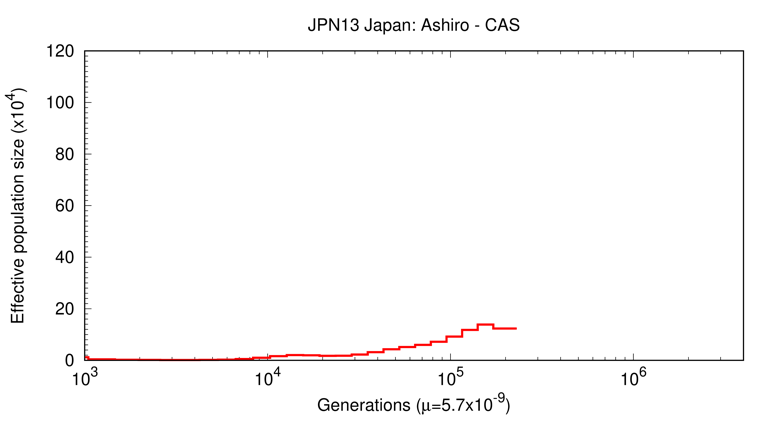

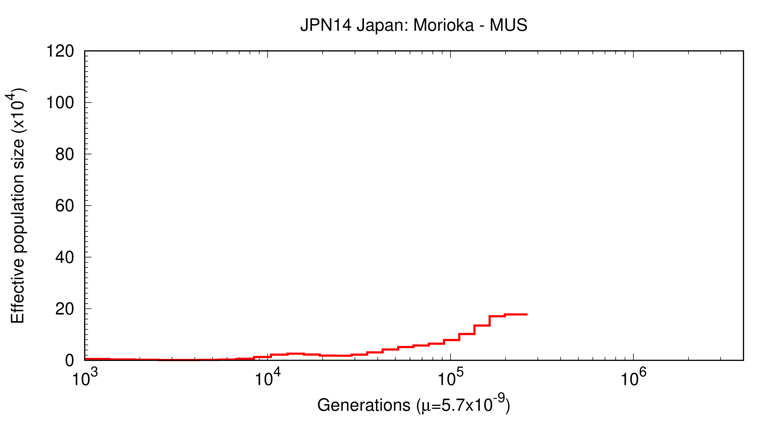

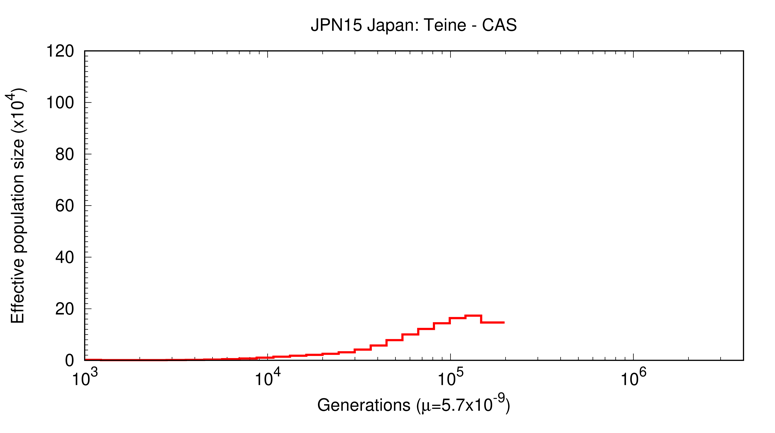

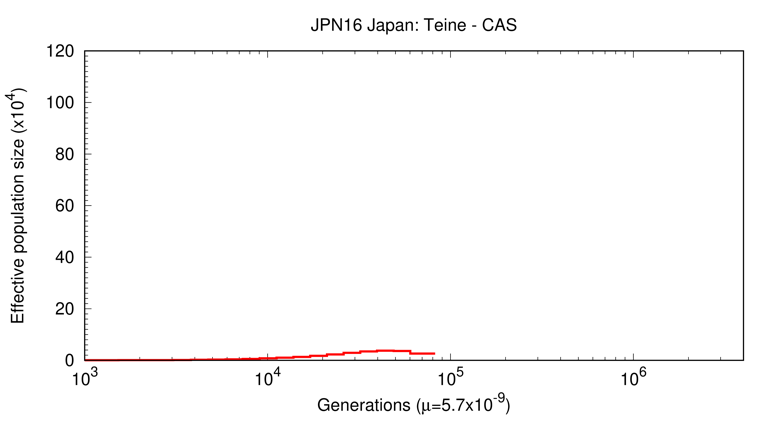

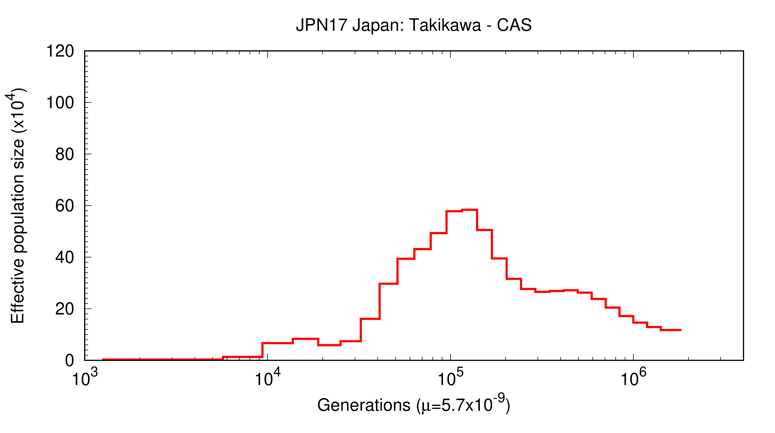

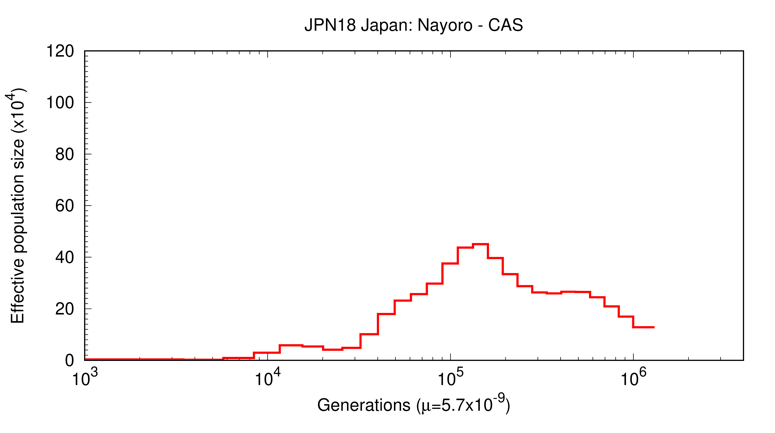

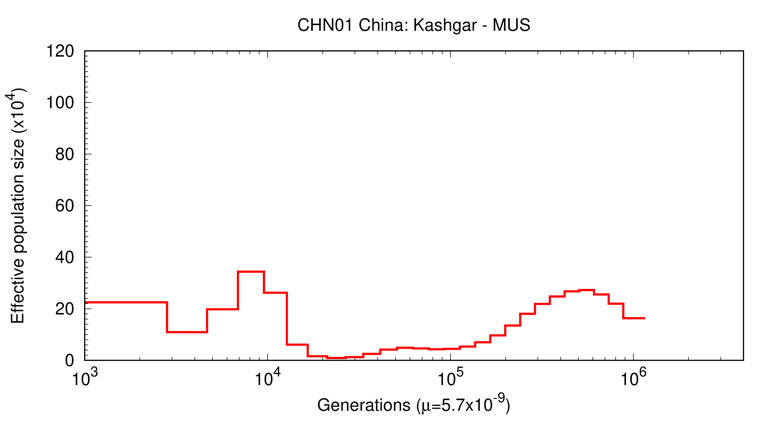

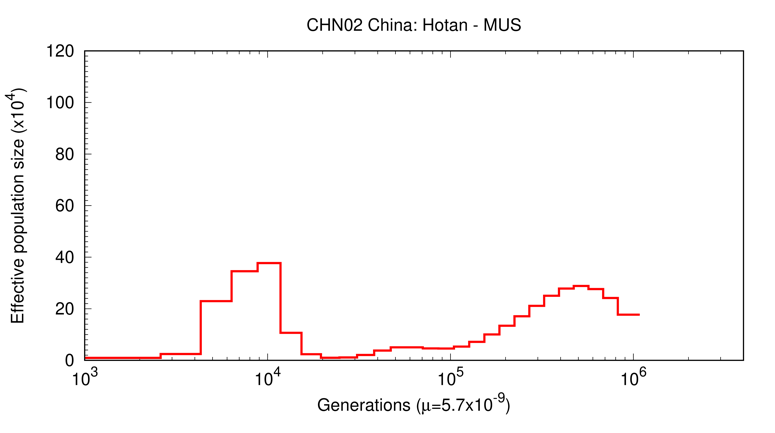

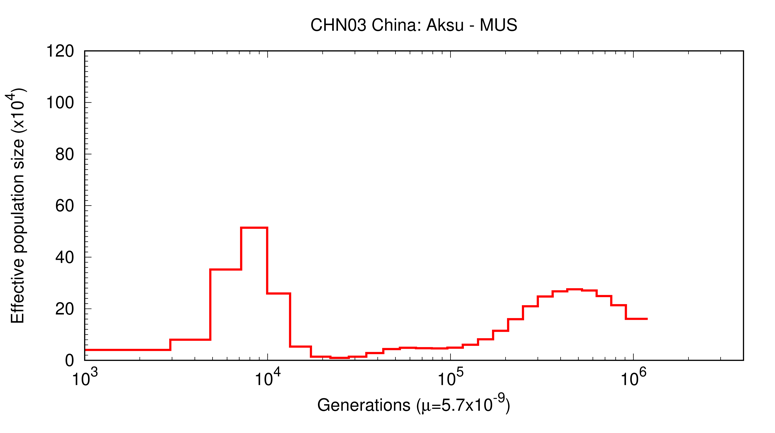

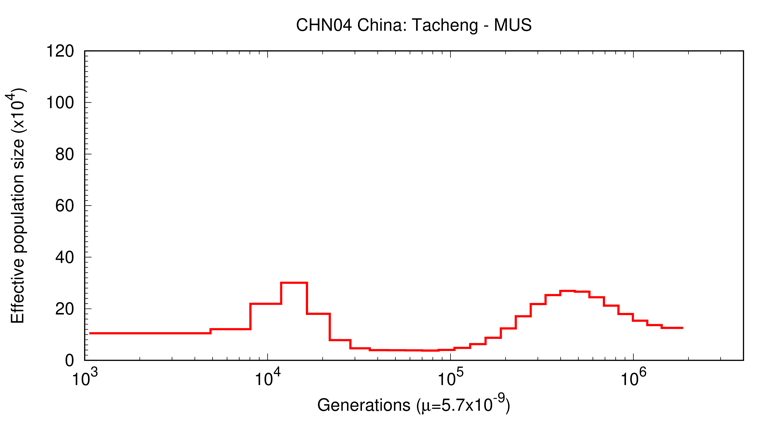

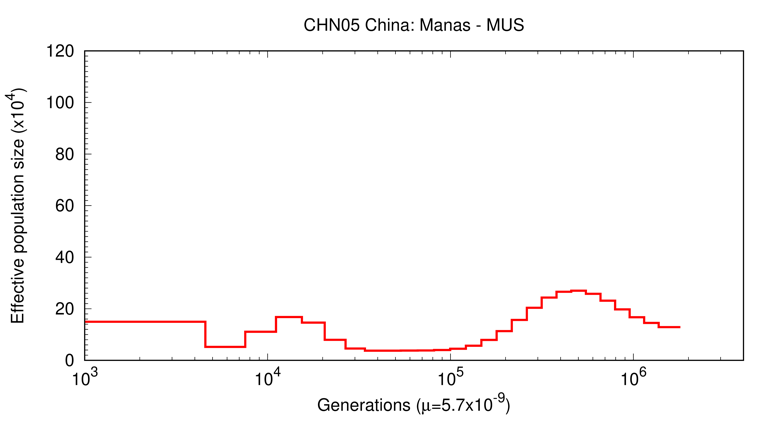

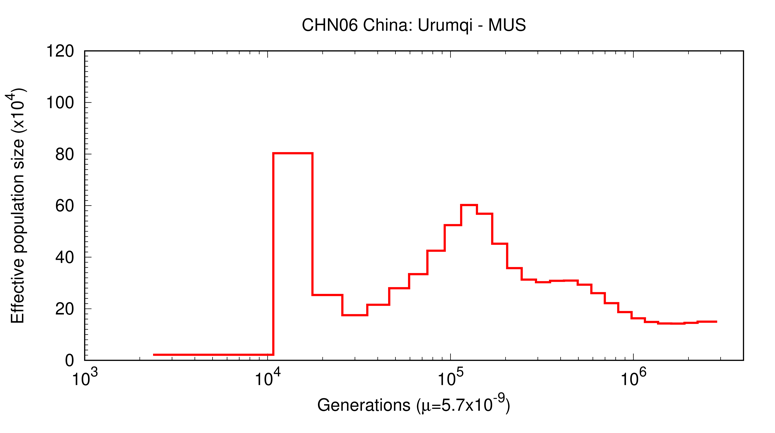

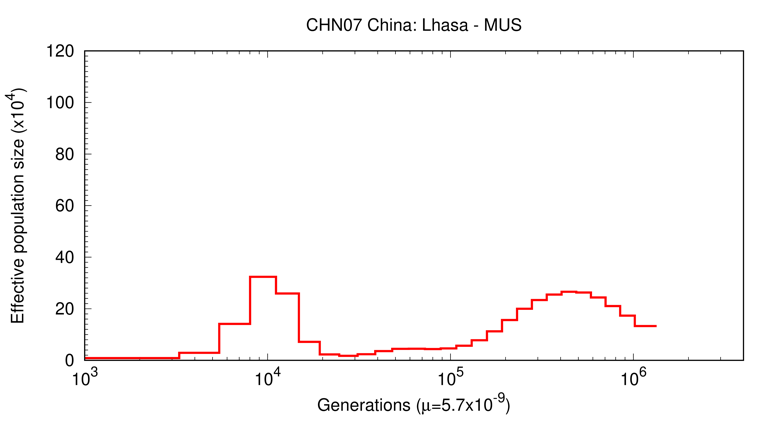

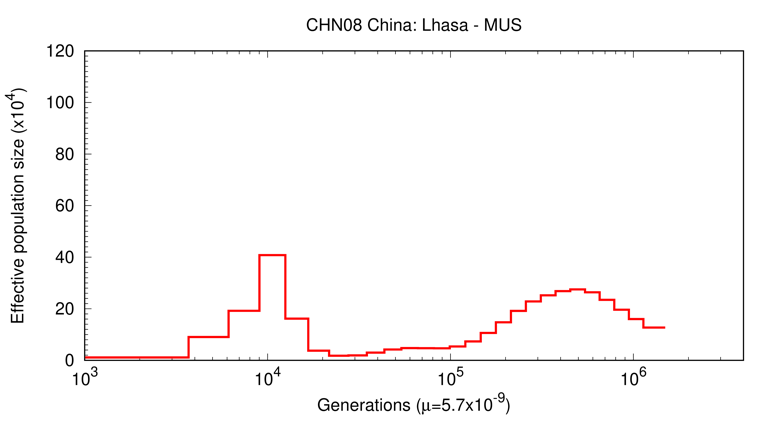

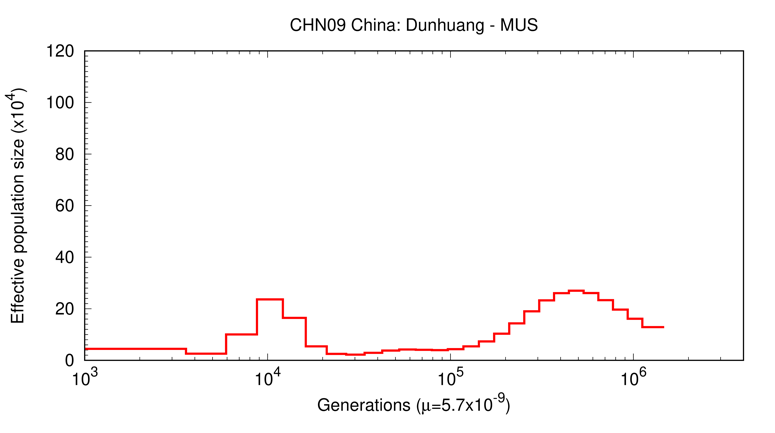

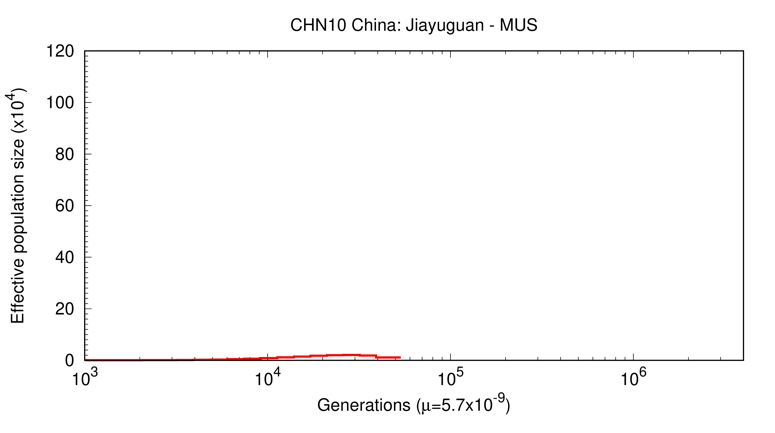

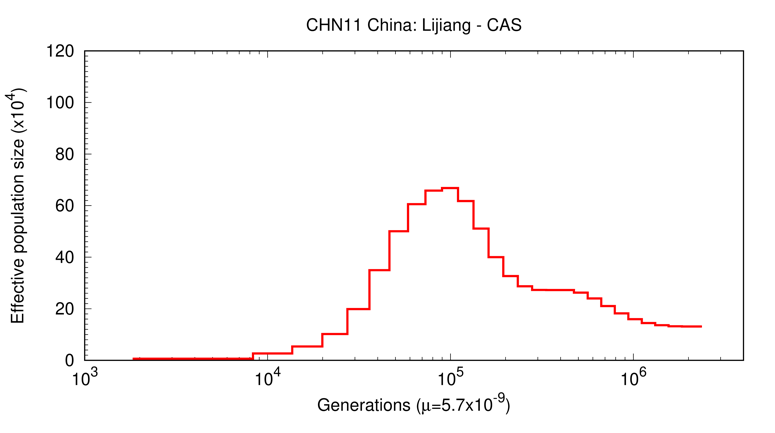

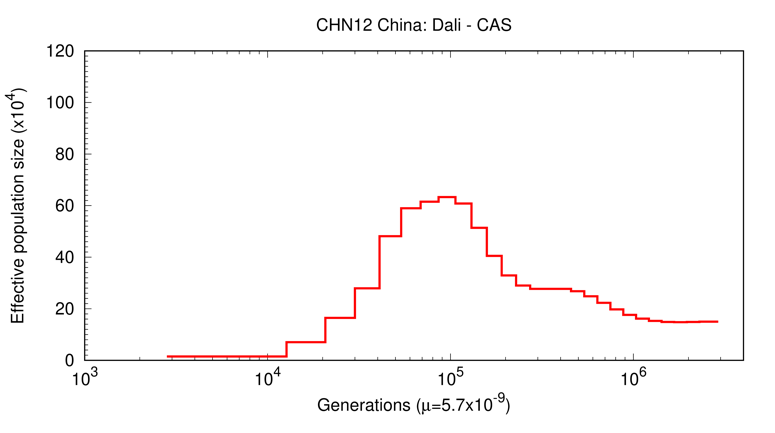

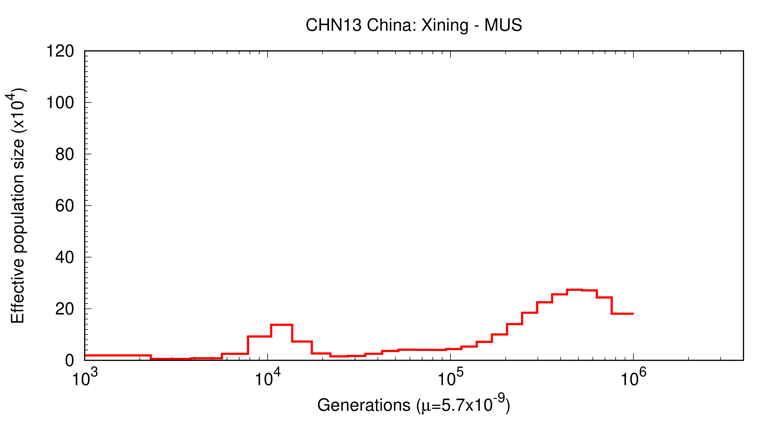

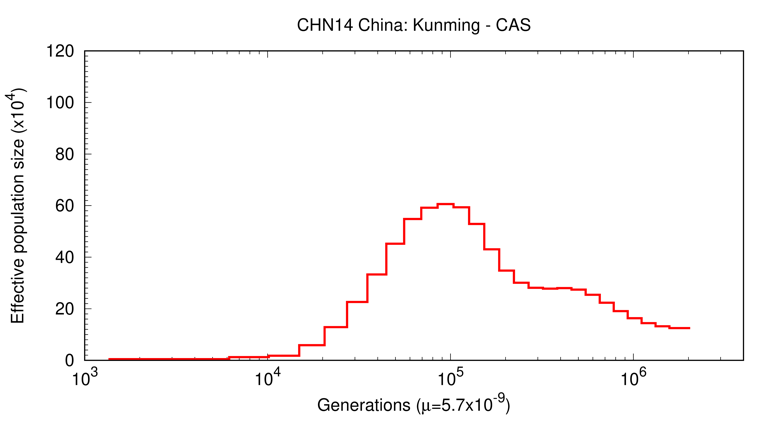

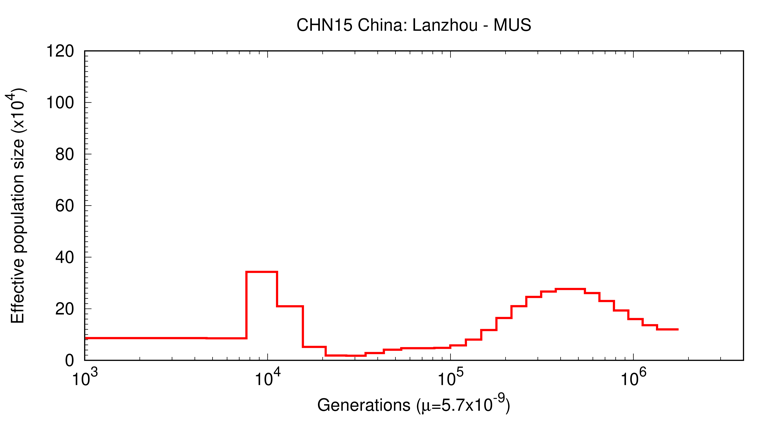

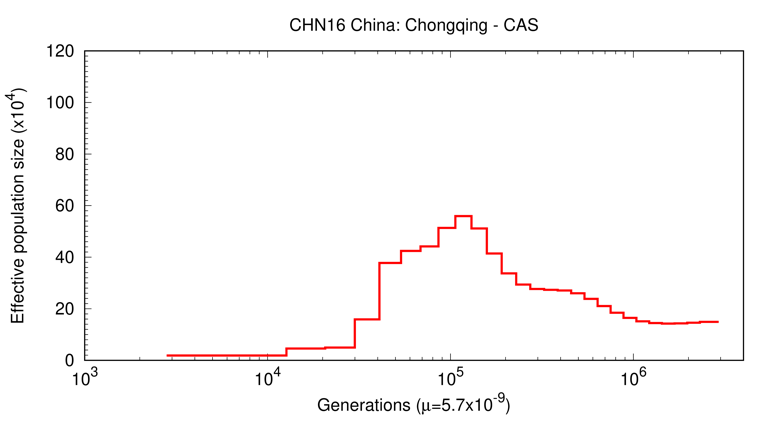

**Supplementary Figure 7**

The PSMC plots of all samples used in this study. The figures are sorted according to the ADMIXTURE plot sample sorting.

**Supplementary Figure 8**

The plot of all pairwise genetic distances between samples (IBS distance). The light-colored cells indicate that the genetic distances between the two samples are close.

**Supplementary Figure 9**

The neighbor-joining tree inferred using all pairwise genetic distance presented in Supplementary Figure 8. The circles next to sample names represent the mtDNA haplogroups of samples.
